# Zero-phase-delay synchrony between interacting neural populations: implications for functional connectivity derived biomarkers

**DOI:** 10.1101/2025.01.04.631256

**Authors:** Chirag Mehra, Ahmad Beyh, Petroula Laiou, Pilar Garces, Emily JH Jones, Luke Mason, Jan Buitelaar, Mark H Johnson, Declan Murphy, Eva Loth, Flavio Dell’Acqua, Joshua B Ewen, Mark P Richardson, Jonathan O’Muircheartaigh

## Abstract

Neural populations synchronise their activity with either zero-phase-delay (activity in interacting regions occurs simultaneously) or a phase-delay (activity in one region follows the other). In electroencephalography and magnetoencephalography functional connectivity analyses, artefactual connectivity can also occur with zero-phase-delay. To minimise artefact, contemporary analyses typically exclude all zero-phase-delay interactions. However, the extent to which ‘true’ interactions are resultingly lost – and the impact this has on the performance of functional connectivity metrics as biomarkers – remains unknown. Here, we show that most cortico-cortical functional connectivity occurs with zero- or near-zero phase-delay, even where such connectivity is unlikely to be artefactual. Including, rather than excluding, zero-phase-delay connectivity increases the reliability, convergence with neurobiology (structure-function concordance, homotopic interhemispheric connectivity, and age-related connectivity changes), and prognostic ability of functional connectivity metrics. We find that excluding zero-phase-delay connections penalises functional connectivity strength between the strongest structurally connected regions: stronger structural connections lead to functional connections with phase-delays closer to zero, mediated by a shorter signal propagation time. Our findings challenge generally accepted assumptions that zero-phase-exclusive methods are superior to zero-phase-inclusive methods.

## Introduction

Populations of neurons produce oscillatory electromagnetic activity (Bishop, 1932; Buzsáki, 2002). Through the precise synchronisation and integration of these oscillations, the brain produces cognitive function, sensory experience and behaviour (Buzsáki, 2006; Lopes da Silva, 2013; Rodriguez et al., 1999a; Singer, 1999; Uhlhaas et al., 2009; Varela et al., 2001; Womelsdorf & Fries, 2007). Studying these interactions, or functional connectivity, has provided insight into several cognitive processes (Dai et al., 2017a; Hampson et al., 2006; Kang et al., 2011) and neuropsychiatric disorders (Du et al., 2018). Indeed, metrics derived from functional connectivity analyses have been proposed as candidate diagnostic or prognostic biomarkers for brain-based conditions (Du et al., 2018; Gao et al., 2020; Gholipour et al., 2022; Gil Ávila et al., 2023; Hong et al., 2020; Plitta et al., 2015).

The averaged oscillatory electromagnetic activity of neural populations can be non-invasively measured by electroencephalography (EEG) and/or magnetoencephalography (MEG; Singer, 1999). Brain-regions are assumed to be functionally connected if the phase and/or amplitude (Engel et al., 2013; Varela et al., 2001) of their oscillations are statistically interdependent (Friston, 2011). Functionally connected brain regions can have perfectly synchronous activity (with zero-phase-delay), or activity in one region may follow the other (producing a phase-delay). However, the extent to which measured zero-phase-delay connectivity reflects ‘true’ versus artefactual connectivity in empirical signals remains unknown.

There is robust evidence demonstrating the presence of ‘true’ zero-phase-delay functional connectivity, replicated across species (Engel et al., 1991; O’Reilly & Elsabbagh, 2021; Vicente et al., 2008; Witham et al., 2007). Spike-train recordings (Engel et al., 1991) have demonstrated that most functional connections between homotopic interhemispheric regions occur with zero-phase-delay. Computational models suggest that “resonance-induced synchrony” (Gollo et al., 2014) facilitates zero-phase-delay connectivity: bidirectional information transfer between regions acts to mutually alter their dynamics into a stable zero-phase-delay pattern. Functions dependent on zero-phase-delay connectivity include maximizing the reliability of information transmission and facilitating spike-time-dependent plasticity (Gollo et al., 2014), coding and binding features of sensory objects (Gray et al., 1989; Rodriguez et al., 1999b; Singer, 1999), and others (Roelfsema et al., 1997; Steinmetz et al., 2000). Evidence from invasive recordings also shows near-zero-phase-delay connectivity (Uhlhaas et al., 2009), describing phase-delays that are much smaller than those arising from axonal conduction delays. An exemplar function of near-zero-phase-delay connectivity is neural encoding by hippocampal place cells (O’Keefe & Recce, 1993).

Artefactual functional connectivity due to signal leakage and volume conduction also occurs with zero-phase-delay (Brunner et al., 2016; Hipp et al., 2012; Nolte et al., 2004; Srinivasan et al., 2007), as spatial blurring in signal leakage (Anzolin et al., 2019; Colclough et al., 2015; Hipp et al., 2012; Srinivasan et al., 2007) and field-spread in volume conduction (Anzolin et al., 2019; Bastos & Schoffelen, 2016; Nolte et al., 2004) occur with zero- or almost-zero-time delay. This artefactual connectivity is unavoidable in EEG and MEG data. It mimics ‘true’ connectivity (Anzolin et al., 2019) and may dominate the pattern of connectivity measurements (Hipp et al., 2012; Srinivasan et al., 2007).

Therefore, contemporary functional connectivity analyses typically exclude zero-phase-delay connections, prioritising the goal of limiting artefactual connectivity due to signal leakage and volume conduction (Colclough et al., 2016; Miljevic et al., 2022). However, the extent to which biologically relevant ‘true’ connectivity is also lost when excluding zero-phase-delay connectivity is unknown. Including versus excluding zero-phase-delay connectivity may substantially alter the conclusions drawn from otherwise identical signals (Cociu et al., 2018), hindering progress in biomarker discovery (for example, O’Reilly et al., 2017). No study, to our knowledge, has systematically assessed the effect of including versus excluding zero-phase-delay connections on the performance of functional connectivity metrics as candidate biomarkers.

Here, using empirical EEG data, we used a novel approach to quantify the proportion of zero- and near-zero phase-delay connectivity when signal leakage artefact was likely negligible. Then, we compared the effects of including versus excluding zero-phase-delay functional connectivity on properties desired in biomarkers: high test-retest reliability (Davis et al., 2020) and convergence with underlying biology (FDA-NIH Biomarker Working Group, 2016). Finally, we compared the performance of functional connectivity metrics derived from zero-phase inclusive versus exclusive methods as prognostic biomarkers for longitudinal changes in cognitive abilities. Thus, our analyses leveraged multiple performance criteria not trivially correlated with each other.

## Methods

### Participants

We included participants from the AIMS-2-TRIALS Longitudinal European Autism Project, described here (Charman et al., 2017; Loth et al., 2017). Written informed consent was obtained from all participants for being included in the study. In this analysis, we included 153 neurotypical participants (33% female), aged 6-31 years, from 5 study sites. For the analyses predicting longitudinal changes in spatial working memory, participants were split into children (6-12 years, n = 41), adolescents (13-17 years, n = 50) and adults (18-31 years, n = 62), Table 1. Included participants needed at least 20 seconds of usable eyes-closed resting-state EEG data, a good quality (defined below) T1-weighted (T1w) magnetic resonance spectroscopy (MRI) scan image, and a full-scale intelligence quotient ≥75. We excluded participants with diagnosed neurodevelopmental conditions, intellectual disability, neurological conditions, mental health conditions (e.g. schizophrenia, bipolar affective disorder) and significant systemic disease (e.g. auto-immune conditions). Intelligence quotient was measured by the Wechsler Abbreviated Scale of Intelligence, Second Edition (Wechsler, 2011).

**Table 1.**
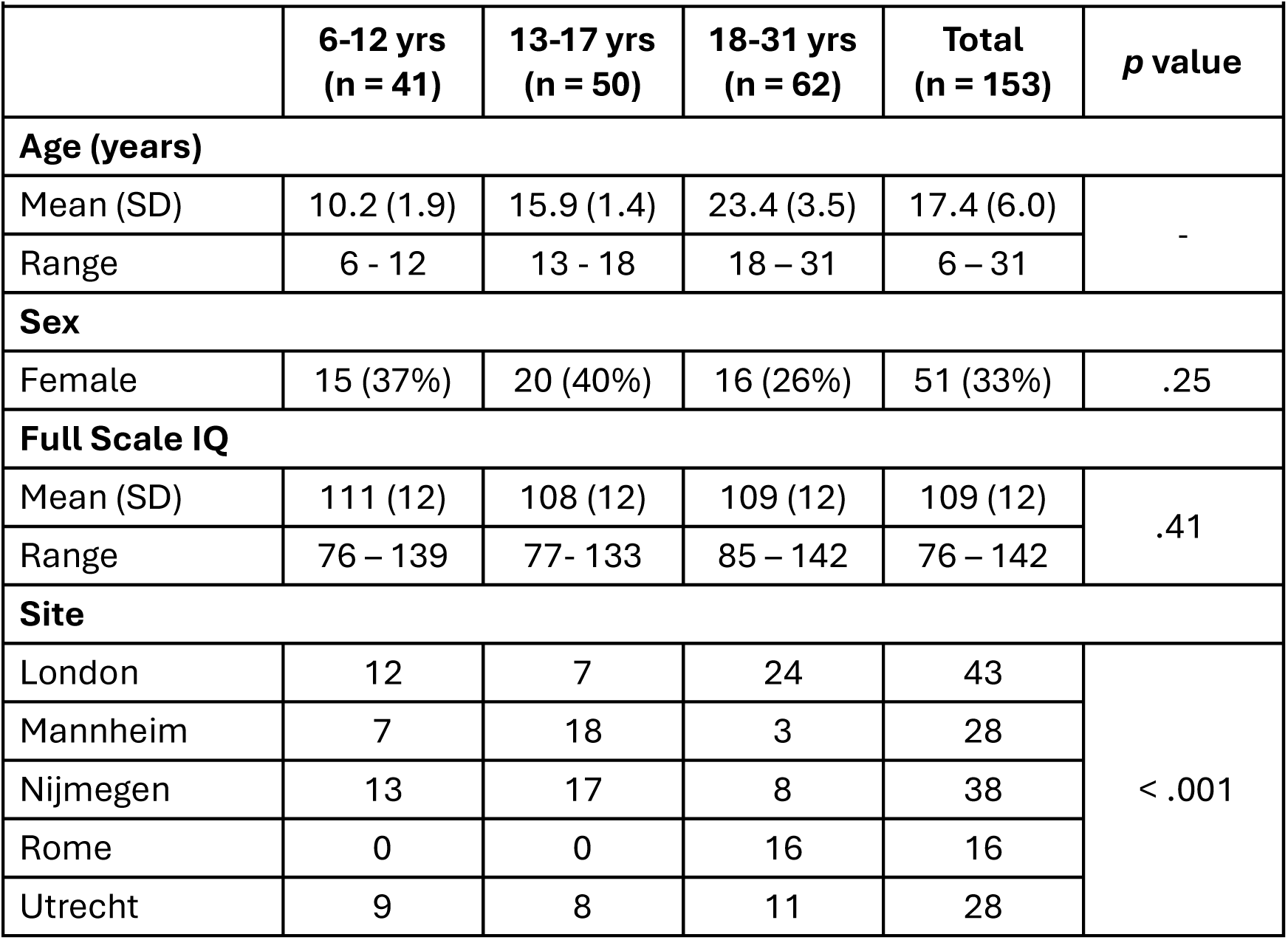
Characteristics of included participants. This study included 153 participants aged between 6 and 31 years. For the analyses predicting longitudinal changes in spatial working memory, participants were grouped by age: children (6-12 years), adolescents (13-17 years), and adults (18-31 years).

### Electroencephalography methods

EEG artefact cleaning and source localisation were performed as per our group’s previous work (Garcés et al., 2022).

#### EEG data acquisition

Four segments of 30 second, eyes-closed EEG recordings were captured. These short epochs were used to maximise comfort for young participants. Data were acquired at five sites: Central Institute of Mental Health (CIMH, Mannheim, Germany), King’s College London (KCL, United Kingdom), University Nijmegen Medical Centre (RUNMC, Netherlands), University Campus BioMedico (UCBM, Rome, Italy) and University Medical Centre Utrecht (UMCU, Netherlands). The following EEG systems were employed: Brainvision (CIMH, KCL, RUNMC), Biosemi (UMCU) and Micromed (UCBM), with sampling frequencies of 5000 Hz (KCL, RUNMC), 2048 Hz (UMCU), 2000 Hz (CIMH) and 256–1000 Hz (UCBM). All sites used 10–20 layout caps, with 60–70 electrodes.

#### EEG artefact cleaning

Only data from the 61 electrodes commonly used across all sites were retained. The following steps were performed: (1) electrode channels with poor data quality were removed, (2) temporal segments with large transient artifacts such as muscle bursts or movements were discarded, (3) independent component analyses were applied using fastICA (Hyvärinen, 1999) to distinguish signal from noise, (4) artefactual independent components were identified and eliminated, (5) and channels discarded in (1) were interpolated. The common average signal across all channels was used as the reference. If more than 10 cannels were eliminated in step 1, the epoch was not included in subsequent analyses.

Prior to source localisation, initial downsampling and bandpass filtering were performed, as per Hyvärinen (1999). Except for signals from UCBM (which were recorded at 256 Hz and not initially downsampled), EEG signals were downsampled to 1000 Hz, with an antialiasing filter (low-pass finite impulse response filter, Kaiser window, cutoff of 300 Hz, transition band width 100 Hz), using EEGlab (Delorme & Makeig, 2004). Then, the entire EEG resting-state recording (all four 30 second epochs and the segments in between them), with 10 seconds of data padding, was bandpass filtered between 1–32 Hz with a finite impulse response filter of order 2000, using a Hamming window, in both forward and backward directions, with detrending, using FieldTrip (Oostenveld et al., 2011).

#### Source Reconstruction

Source reconstruction was performed using individual-participant T1-weighted MRI images (acquisition parameters detailed below). T1-weighted MRIs were segmented with Statistical Parametric Mapping 12 (SPM12) software (Penny W et al., 2006) into grey matter, white matter, cerebrospinal fluid, bone, soft tissue and air. These probabilistic images were then smoothed (5mm full width half maximum), thresholded and resliced to produce binary masks of 2mm × 2mm × 2mm resolution for three tissue types: brain (including grey matter, white matter and cerebrospinal fluid), skull and scalp. These binary masks were transformed to hexahedral meshes with FieldTrip (Oostenveld et al., 2011). All three considered tissue types were assumed to have homogeneous and isotropic conductivity: 330 ms/m for the brain and scalp (Gabriel et al., 1996), and an age-dependent skull conductivity of 3.958 + 62.77 x *e*^− 0.2404 x age in years^ ms/m, in line with the BESA (*BESA® | Brain Electrical Source Analysis: Home*, 1995) recommended conductivity ratios. Segmentations were visually inspected. The forward model was derived with FieldTrip and SimBio (Vorwerk et al., 2018). For the inverse model, 1200 source locations of interest were defined in grey matter in Montreal Neurological Institute (MNI) space, following a 3D cubic diamond grid. Source positions were transformed from MNI space to each subject’s individual space with a nonlinear transformation using SPM12. Electrode positions were determined by transforming standard MNI positions to subject-space with the same transformation, then projecting to the scalp surface. Source time series were estimated with linearly constrained minimum variance beamformer (Anzolin et al., 2019), using a regularization of 5% of the average trace of the covariance matrix. The 1200 sources were parcelled into 68 cortical regions of interest, as per the Desikan-Killiany atlas (Desikan et al., 2006). Each region was comprised of between 1 and 67 sources (median = 13). When there was more than one source per region, the representative time series for a given region was defined as the first principal component of all the source time series in it.

### Cropping, down-sampling and filtering

We chose 20 second eyes-closed resting-state EEG epochs (from a maximum segment length of 30 seconds) to produce functional connectivity adjacency matrices. This epoch length was chosen as a compromise between maximising the number of included participants and maximising network stability (Fraschini et al., 2016; Pullon et al., 2020).

We then further down-sampled the signals to 250Hz. Data quality checks were subsequently performed.

We treated the whole 20 second epoch as a single window. Producing multiple windows has the advantage of better capturing moment-to-moment connectivity. However, longer window-length better captures the ‘ground truth’ (Pullon et al., 2020). Additionally, longer windows are needed to capture weaker connections (Pullon et al., 2020) due to the noise inherent to EEG signals.

The signals were then filtered into the conventional frequency bands: delta (1-4 Hz), theta (4-8 Hz), alpha (8-13 Hz), low-beta (13-20 Hz) and high-beta (20-32 Hz). High-pass followed by low-pass filters were used in sequence, using two-pass (using the MATLAB function filtfilt), finite impulse response filters, of order 250. To reduce signal distortion, data padding of 500 samples (2 seconds) was used during filtering and while performing Hilbert transformations.

### Quantifying phase-delays

To quantify the phase-delay between two signals *x*_*i*_ (*t*) and *x*_*j*_(*t*), *t* = 1,…., *N* where *N* is the number of points in an EEG epoch, we first applied the Hilbert transform to obtain the instantaneous phases *θ*_*i*_(*t*), *θ*_*j*_(*t*) for the signals *x*_*i*_ (*t*) and *x*_*j*_(*t*) respectively. Then, the phase-delay distribution Δ*θ*(*t*) = *θ*_*i*_(*t*) − *θ*_*j*_(*t*) was projected into the unit circle. The instantaneous phase-delays across samples in an epoch were averaged to produce the mean phase-delay throughout the epoch:

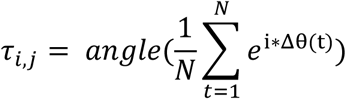

### Zero- and ±π-phase-delay functional connectivity

Note that ‘true’ ±π-phase-delay functional connectivity also occurs between region-pairs where ‘true’ zero-phase-delay connectivity is prevalent (Ni et al., 2009; O’Reilly & Elsabbagh, 2021; Palmer et al., 2012), and has important functions (Hilgetag et al., 2001; Ni et al., 2009; O’Reilly & Elsabbagh, 2021; Palmer et al., 2012). As ±π-phase-delay functional connectivity is also excluded by zero-phase-exclusive functional connectivity methods, for brevity, zero-phase-delay and π-phase-delay are subsequently referred to together as ‘zero-phase-delay.’

### Investigating where the effects of signal leakage are likely negligible

We graphically examined the relationship between the proportion of zero and near-zero phase-delay functional connectivity and Euclidean distance between region-pairs. Artefactual connectivity due to signal leakage occurs at zero-phase-delay (Gohel et al., 2017; Hipp et al., 2012) and its influence is a function of Euclidean distance between sources (Bastos & Schoffelen, 2016; Gohel et al., 2017; He et al., 2019; Hipp et al., 2012). Therefore, we explored if there were Euclidean distances between brain regions at which the proportion of zero- and near-zero phase-delay connectivity did not vary as a function of Euclidean distance. Such a phenomena would suggest that artefactual connectivity due to signal leakage is likely negligible at these Euclidean distances.

Euclidean distance between region-pairs was measured from the centroids of the regions, obtained from the FreeSurfer surface parcellation, measured per participant using individual T1w MRI images.

In the absence of an established definition of ‘near-zero-phase-delay’ functional connectivity, we defined near-zero-phase-delay as within < | 0.3 | radians from 0 or ±π radians. These phase-delays comprise approximately 19% of possible phase-delays in a unit circle (between −π and π). For comparison, the phase delay caused by signal transmission delays along a (white matter) streamline of length 128mm (the median streamline length in our dataset), assuming a white matter transmission speed of 4.5m/s (van Blooijs et al., 2023) and a synaptic transmission delay of 0.75 milliseconds (Katz & Miledi, 1965), for oscillations at 10 Hz, is approximately 1.83 radians. Note that by design, all zero-phase-exclusive methods alter the functional connectivity value based on how close phase-delay is to 0 or π. Connectivity strength values are only unaltered by zero-phase-exclusive methods at phase delays of ±π/2 radians, and approach zero as phase-delay approaches 0 or ±π. At phase delays of 0.3 radians, the measured connectivity strength of ‘true’ (and artefactual) functional connectivity is decreased by 70% using the wPLI and Img COH compared to when using zero-phase-inclusive methods.

## Functional Connectivity Methods

We limited our selection of methods for comparison to those that characterise functional connectivity based on phase relationships, amplitude relationships, or a combination thereof; do not make assumptions about directionality of connectivity based on the sign of phase difference (Stam & van Straaten, 2012); are bivariate; and are commonly used. Thus, we selected phase locking value (PLV; Lachaux et al., 1999), amplitude envelope correlation (AEC; Bruns et al., 2000) and coherence (Adey et al., 1967) as zero-phase-delay inclusive methods. We selected their methodologically corresponding zero-phase-delay exclusive methods: weighted phase lag index (wPLI, Vinck et al., 2011), orthogonalized amplitude envelope correlation (Orth AEC; Hipp et al., 2012) and the imaginary part of coherency (Img COH; Nolte et al., 2004), respectively.

### Coherence and imaginary part of coherency

Coherence measures the consistency of relative amplitude and phase between pairs of signals at a given frequency.

Coherence is the absolute value of coherency, C*_ij_*(f), between signals *i* and *j*, defined as:

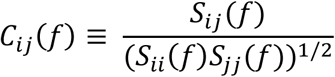

Where *S_ij_(f)* is the cross spectral density of signals *i* and *j, S_ii_(f)* is the power spectral density of signal *i,* and *S_jj_(f)* is the power spectral density of signal *j*. We used the Welch method to calculate power spectral density, as it has the advantage of handling signal leakage effects well (Welch, 1967).

Therefore, coherence Coh*_ij_*(f), is:

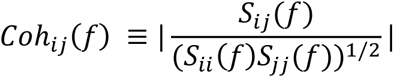

The imaginary part of coherency only considers the imaginary component of the complex-valued cross spectral density:

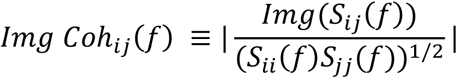

### Phase Locking value

Phase-based connectivity methods assume that two brain regions are connected if the rhythm of their oscillations are similar, largely independent of amplitude relationships.

The phase locking value between the signals *x*_*i*_ (*t*) and *x*_*j*_(*t*), *t* = 1,…., *N* where *N* is the number of points in an epoch is defined as:

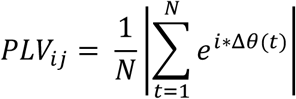

Where Δ*θ*(*t*) is the difference of the instantaneous phases *θ*_*i*_ (*t*), *θ*_*j*_(*t*) of the signals *x*_*i*_(*t*), *x*_*j*_(*j*). Here, we did not perform the permutation testing that is typical for the PLV (Lachaux et al., 1999), to enable comparisons with the other structural and functional connectivity methods used in this analysis.

### Weighted Phase Lag Index

The weighted phase lag index is derived by multiplying the phase lag index value by a multiplier based on the amplitude of the phase-difference.

The weighted phase lag index between two signals *i* and *j* is defined as:

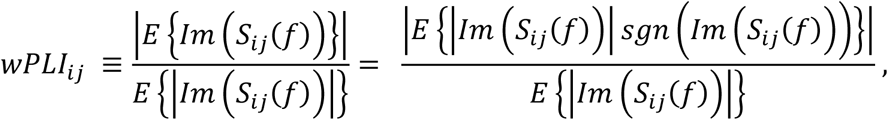

where *S*_*ij*_ is the cross spectrum between the signals *i* and *j*.

### Amplitude envelope correlation and orthogonalized amplitude envelope correlation

The amplitude envelope correlation (AEC) detects amplitude relationships between signals, independent of phase relationships. The AEC between region-pairs *i* and *j* is the magnitude of the Pearson correlation coefficient between the absolute values of Hilbert transformed signal *i* and the absolute values of Hilbert transformed signal *j*.

In the orthogonalized AEC, pairs of signals are orthogonalized with respect to each other prior to carrying out the AEC calculation. Orthogonalization was performed (on a per-sample basis) in two directions (*i* to *j*, *j* to *i*), producing *i*_⊥_*_j_* and *j*_⊥_*_i_* timeseries, as per Hipp and colleagues (2012). *i*_⊥_*_j_*, for example, was derived by:

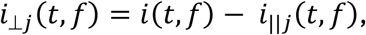

when

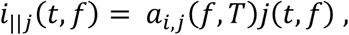

Where *a_i,j_* is the regression coefficient obtained by ordinary least squares describing the linear relationship between *i* and *j* throughout the epoch (t = 20). The orthogonalized time series were then squared and log transformed, to make the time series more normal, as per Hipp and colleagues (2012). Then, power envelope correlations were computed between *i* and *j*_⊥_*_i_*, and *j* and *i*_⊥_*_j_* – the mean of the two correlation values was taken to be the undirected orthogonalized connectivity value between *i* and *j*.

### Test-retest reliability

We quantified the intrasession test-retest reliability between two epochs recorded between 1-3 minutes of each other (n = 99). Intrasession test-retest reliability was used to reduce the confounding effects of circadian rhythm (Fafrowicz et al., 2019) and caffeine (Haller et al., 2013) on functional connectivity. Two measures of test-retest reliability were used: absolute agreement and consistency. Edgewise absolute agreement was quantified using the interclass correlation coefficient (ICC (2,1)). ICC values were interpreted as per Koo & Li (2016): 0.9 −1.0 indicate excellent reliability, 0.75-0.9 indicate good reliability, 0.5-0.75 indicate moderate reliability and 0.0-0.5 indicate poor reliability. Negative values were given a value of 0. We quantified the consistency of the whole adjacency matrix using Spearman’s correlation coefficient, as per van der Velde and colleagues (2019).

### Structural Connectivity

Structural connectivity between 68 grey matter regions, as parcellated by the Desikan-Killiany atlas (Desikan et al., 2006), was quantified by using spherical deconvolution diffusion weighted imaging. Diffusion imaging data was available from 2 of 5 study sites (KCL and Mannheim), comprising 50 participants (Table 2a). Spherical deconvolution diffusion imaging was used to overcome the crossing fibres problem that is a key limitation of traditional diffusion tensor imaging (Dell’Acqua et al., 2013). Connectivity strength between two regions was based on the streamline count between them, normalised to the value of 1, as per Chu and colleagues (2015).

**Table 2a.** demographics of participants who had both EEG and diffusion weighted imaging data. FSIQ: full scale intellectual quotient

| Participants | KCL n = 28 | Mannheim n = 22 |
| --- | --- | --- |
| Mean Age (SD) | 23 (4.7) | 15 (2.9) |
| Age Range | 15 - 31 | 8 - 19 |
| Mean FSIQ (SD) | 109 (13.3) | 110 (13.3) |
| FSIQ Range | 85 – 142 | 77 – 133 |

### T1-weighted cortical reconstruction using FreeSurfer

Signals from diffusion imaging were projected on to the cortical surface derived from each participant’s T1w MRI image. As previously done by our group (Pretzsch et al., 2022), all T1 scans were manually inspected; those with visible anomalies or significant movement artefacts were excluded. For each T1w MRI, models of the cortical surface using FreeSurfer v6.0 (https://surfer.nmr.mgh.harvard.edu/) were computed. A detailed description of these well-validated and automated procedures is provided by Fischl (2012). In brief, FreeSurfer performs intensity normalization, skull stripping, and an image segmentation using a connected-components algorithm. Next, it generates a filled white matter volume for each hemisphere and, by fitting a deformable template, a surface tessellation for each of these volumes. For each subject, this yields a mesh (of triangular elements) for the inner (white-matter) and outer (pial) cortical surface consisting of ∼150k vertices (points in each triangular element) per hemisphere. Each of these reconstructed surfaces was inspected visually for reconstruction errors by three independent and experienced researchers. Scans were either accepted or manually edited after automatic pre-processing. Manually edited images were iteratively re-pre-processed and visually re-assessed.

### Diffusion MRI pre-processing

Diffusion MRI acquisition parameters are detailed in Table 2b. Tools from Connectome Workbench (https://www.humanconnectome.org/software) were used to re-tessellate the T1w surfaces of each participant to the common tessellation of the ‘32k_FS_LR’ template. This was done to reduce the number of vertices in each surface to 32,492 points, reducing the computational burden of the cortical projection step.

**Table 2b.**
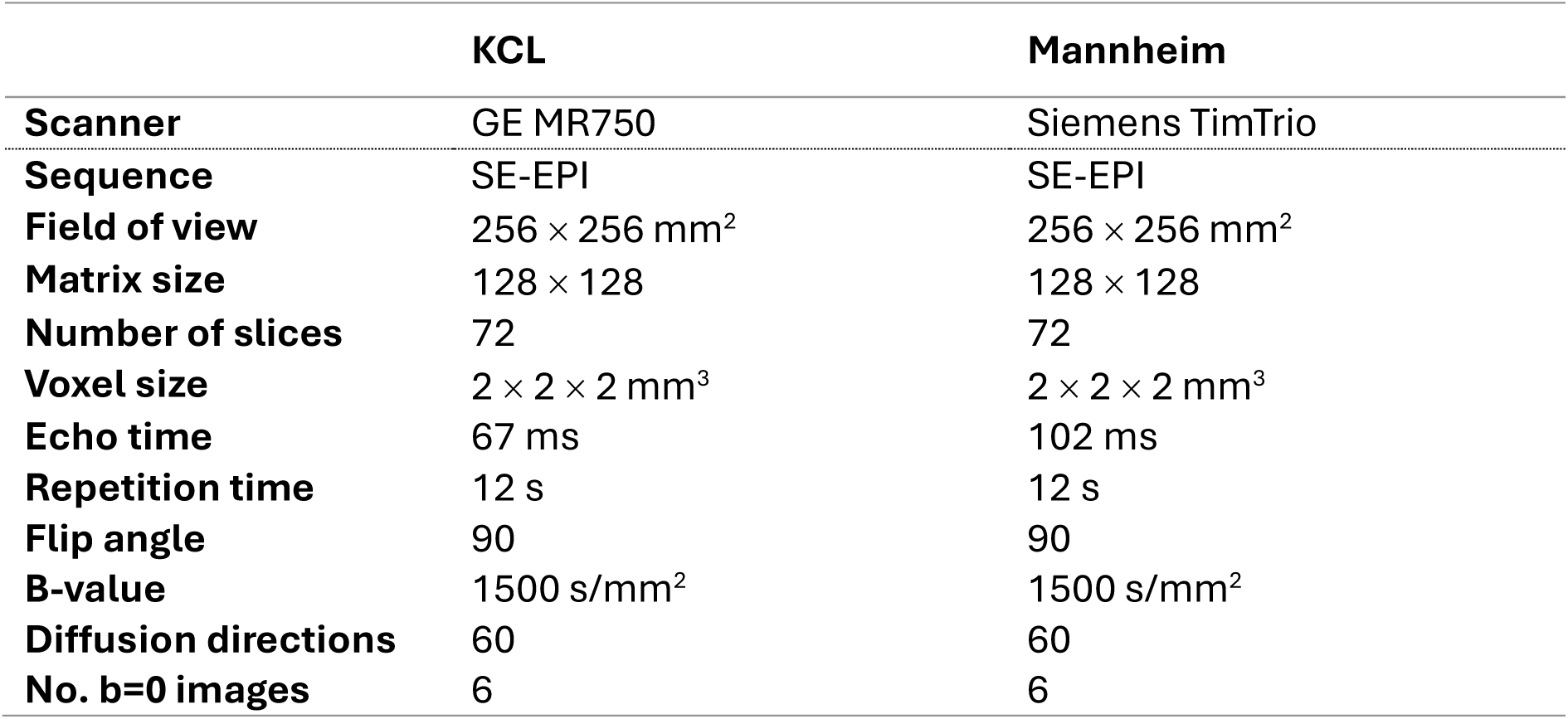
Diffusion MRI acquisition parameters.

The diffusion series of each participant was corrected for thermal noise (Veraart et al., 2016) and Gibbs ringing artefacts (Kellner et al., 2016). Due to the lack of reversed phase encoding images, the *SynB0* framework (Schilling et al., 2019) was used to generate undistorted b=0 EPIs and these were used to estimate the susceptibility distortion field using *topup* (Andersson et al., 2003). Next, *eddy* was used to correct the diffusion series for artefacts caused by frame-wise motion and eddy currents (Andersson & Sotiropoulos, 2016), intra-frame motion (Andersson et al., 2017), and signal dropout (Andersson et al., 2016), as well as susceptibility distortions using the field generated by *topup*. The anisotropic map (Dell’Acqua et al., 2014) was generated from the corrected data and used to calculate a rigid-body registration of the diffusion data to the subject’s T1w structural image in FLIRT (Jenkinson et al., 2002), and this transformation was applied to the entire diffusion series with a spline interpolation to align it with the T1w image; the bvecs were rotated with the same affine matrix.

### Tractography and structural connectivity

Diffusion modelling was performed using the damped Richardson-Lucy spherical deconvolution algorithm (Dell’Acqua et al., 2010) in *StarTrack* (https://mr-startrack.com), with the following parameters: fibre response *α* = 1.5; number of iterations = 300; amplitude threshold *η* = 0.0015; geometric regularisation *ν* = 16. Fibre tracking was performed with the Euler-type algorithm according to the following parameters: minimum HMOA threshold = 0.002; maximum angle threshold = 45°; step size = 1 mm; minimum fibre length = 20 mm; maximum fibre length = 300 mm.

Each participant’s resulting tractogram was then converted to a connectivity matrix, as previously done by our group (Beyh et al., 2022). In brief, each streamline was projected to its nearest neighbouring vertex along the grey-white matter boundary surface (obtained from *FreeSurfer*) with a maximum distance of 3 mm. This resulted in a vertex-wise matrix, which was collapsed into a smaller matrix based on the Desikan-Killiany cortical parcellation (Desikan et al., 2006). Several parcel-wise connectivity matrices were produced with the following edge weights: streamline count; log-transformed streamline count; hindrance modulated orientational anisotropy (HMOA; Dell’Acqua et al., 2013), a tract-specific measure of white matter microstructure; and median fibre length.

### Structure-Function Concordance

Structure-function concordance was calculated on a per-participant basis (n = 50), using the spearman correlation coefficient.

### Predicting age

We assessed the ability of mean strength derived from each functional connectivity method to predict current age over and above head circumference. First, we calculated the R^2^ from 10-fold cross-validation using head circumference as the only predictor of current age (model 1). Then, we added mean strength this model (model 2). We used the change in R^2^ (model 2 – model 1) as the quantity of interest. 147 participants had head circumference data. We excluded two outlier values (90cm from a 22-year-old and 66cm from a 13-year-old) as these likely reflected measurement inaccuracies.

We assessed if model 2 – model 1 cross-validation R^2^ between methodologically corresponding zero-phase inclusive and exclusive methods (e.g. PLI versus wPLI) were significantly different using permutation testing. We created a null distribution by permuting age 1000 times for each method-pair in each frequency band.

### Prognostic ability for longitudinal changes in spatial working memory

Spatial working memory was measured using the ‘Find the Phone’ Task (Owen et al., 1990; Sjöwall et al., 2013). In this task, multiple telephones were shown on a computer screen and ‘rang’ one at a time. Through trial and error, participants had to guess which phone was ringing. Participants were instructed that each phone would only ring once within a trial. The task was to remember which telephone(s) had already rung to avoid re-selecting that phone. Instances where a participant selected a phone that had already rung, or had previously been incorrectly selected, were counted as errors. The metric of interest was ‘total participant errors.’

For each functional connectivity method, we used the weighted, normalised pathlength in the theta band as the candidate prognostic biomarker. The theta band has been robustly associated with working memory (Bauer et al., 2021a; Dai et al., 2017a; Fell & Axmacher, 2011; Hyman et al., 2010; Jones & Wilson, 2005; Siapas et al., 2005). We only looked at connectivity between region-pairs with a strong evidence-base of being implicated in spatial working memory, including regions in the prefrontal cortex (MacKey & Curtis, 2017; Siapas et al., 2005; Wager & Smith, 2003; Wirt & Hyman, 2017), the posterior parietal cortex (Almeida et al., 2015; MacKey & Curtis, 2017), the anterior cingulate cortex (Wirt & Hyman, 2017), and the entorhinal cortex (Almeida et al., 2015; Salimi et al., 2021), as defined by the Desikan-Killiany atlas (Desikan et al., 2006). In total, there were 14 regions of interest in each hemisphere. Shorter weighted, normalised pathlengths reflect higher network efficiency. Global network efficiency and pathlength have been associated with working memory performance (Langer et al., 2013; Stanley et al., 2015).

To calculate weighted, normalised pathlength, we first created weighted, undirected functional connectivity adjacency matrices (networks) with 28 nodes. One matrix was created per functional connectivity method. Each node corresponded to a cortical region of the Desikan-Killiany atlas implicated in spatial working memory, while the weighted edges were the functional connectivity strengths between pairs of nodes. Next, using the MATLAB brain connectivity toolbox (Rubinov et al., 2009), we inverted network edges to lengths (edges with larger weights produce shorter lengths). Then, we computed the *shortest possible* distance between all pairs of nodes. Finally, we calculated the average shortest distance between all pair of nodes to produce the weighted normalised pathlength. Formally, the shortest weighted path length *d*_*ij*_ between any two nodes, e.g. *i* and *j* is defined as:

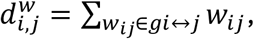

where *gi* ↔ *j* is the shortest weighted path between nodes *i* and *j*.

The weighted, average shortest pathlength (*L*) between node *i* and all other nodes of the network is defined as:

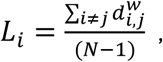

where *N* is the number of nodes (28 in this analysis) and *d*_*i*,*j*_ is the shortest path length between nodes *i* and *j*, when *j* is any node that is not *i*.

To calculate the normalised weighted pathlength, we used a normalisation process by creating 500 random networks by randomising the empirical network, while preserving the degree and strength distributions: each edge was rewired 5 times, with edge weights randomised every 5th step. The weighted pathlength of each random network was calculated, and then averaged across each of the 500 random networks to produce *L*_*rand*_. The normalised average shortest pathlength of node *i* was calculated as:

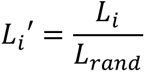

*L*_*i*_′ at each node was averaged across 28 nodes to produce the mean normalised, weighted pathlength (referred to as normalised, weighted pathlength).

Using general linear models, we compared the ability of normalised, weighted pathlength produced from each functional connectivity method to predict changes in total participant errors between the time the EEG was recorded and a time 12-30 months in the future (mean = 19 months, SD = 3.5 months).

### Penalisation Scale

We created a penalisation scale to quantify how connectivity values would be altered by zero-phase-exclusive methods. The scale was between 0 and 1, where the most penalizable phase differences (0 and ±π) were given a value of 1 and least penalizable phase differences (±π/2) were given a value of 0. Penalisation scale values were calculated as follows:

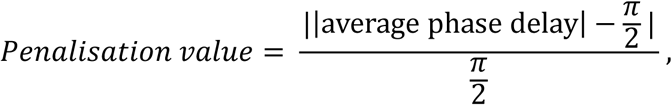

where the average absolute phase delay was limited between 0 and +π. To examine if stronger structural connections were more likely to be associated with more penalizable phase-delays, we spearman correlated the penalisation values with the structural connectivity matrix on a per-participant basis. This was only done for region-pairs between which there were 1 or more streamlines.

### Proxy of signal transmission time

We used a proxy of signal transmission time through white matter tracts. We divided the median length (distance) of white matter tracts between region-pairs by a proxy of transmission speed. The median hindrance modulated orientational anisotropy (HMOA) was used a proxy of transmission speed, as white matter microstructural integrity can reflect myelination (Dell’Acqua et al., 2013) and correlates with nerve conduction velocity (Heckel et al., 2015; Wang et al., 2022). Additionally, the transmission latencies of evoked potentials (an established method of approximating transmission speed; van Blooijs et al., 2023) negatively correlate with white matter microstructure across studies (Evstigneev et al., 2013; Whitford et al., 2011).

### Bootstrapping for mediation effects

We performed a mediation analysis to investigate if the association between structural connectivity strength and the penalisability of the phase-difference of functional connectivity was mediated by a proxy for white matter signal transmission time. We tested the significance of the indirect effect using bootstrapping procedures.

Unstandardized indirect effects were computed for each of 1000 bootstrapped samples, and the 95% confidence interval was computed by determining the indirect effects at the 2.5th and 97.5th percentiles.

## Results

### Most functional connectivity occurs with zero- and near-zero phase-delay, even when artefactual connectivity is likely negligible

We quantified the distribution of the average phase-delay of functional connectivity between brain region-pairs (Fig. 1b). Between all region-pairs across all participants, most measured functional connectivity occurred with mean phase-delays of either zero-/near-zero or π-/near-π radians. Across frequency bands, alpha had the lowest proportion of zero- and near-zero phase delay connectivity (52% of all connections), while high-beta had the highest proportion (69% of all connections), Fig. 1b.

**Fig. 1:**
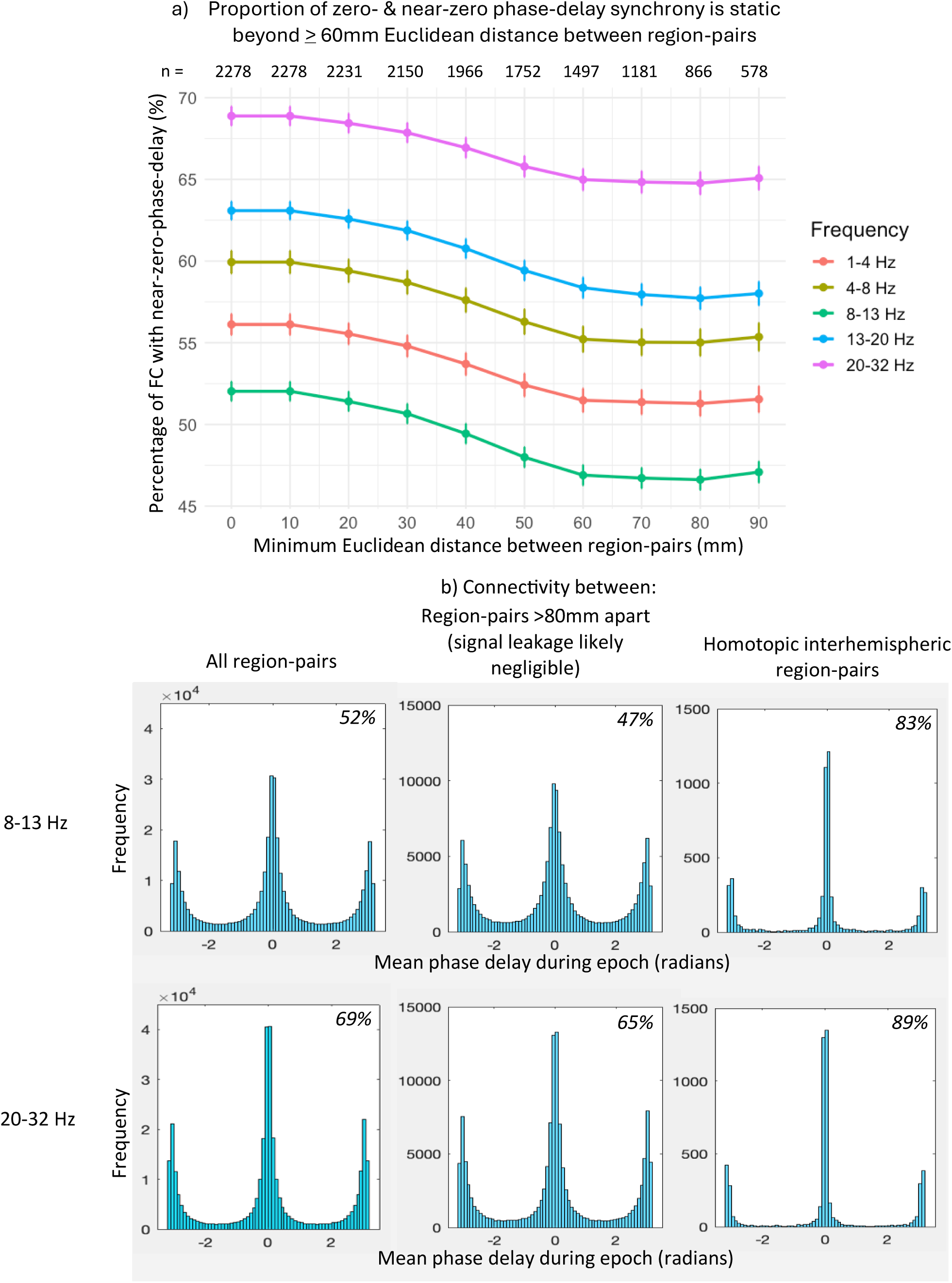
Quantifying zero- and near-zero phase-delay functional connectivity. **a)** Where the percentage of zero- and near-zero phase delay connectivity does not vary as a function of Euclidean distance, artefactual connectivity due to signal leakage is likely to be negligible. This was found to be at Euclidean distances of ≥ 60mm for all frequency bands. The number of pairs of regions per person at each minimum distance is listed above the graph. **b)** Histograms of the average phase-delay of functional connectivity during an epoch between all region-pairs, region-pairs > 80mm Euclidean distance apart (where signal leakage is likely negligible) and between homotopic interhemispheric region-pairs. n = 153. 8-13 Hz and 20-32 Hz are the frequency bands with the lowest and highest near-zero phase delay connectivity, respectively. *Italics*: the proportion of zero- (and ±π) and near-zero (and near ±π) connectivity. FC = functional connectivity.

In source reconstructed data, signal leakage (and not volume conduction) is thought to be the primary contributor of artefactual zero-phase-delay connectivity (He et al., 2019). To quantify the proportion of zero- and near-zero phase-delay functional connectivity when artefactual connectivity due to signal leakage was likely negligible, we used a novel approach. We leveraged the fact that artefactual connectivity due to signal leakage occurs at zero-phase-delay (Gohel et al., 2017; Hipp et al., 2012) and decreases with increasing Euclidean distance between sources (Bastos & Schoffelen, 2016; Gohel et al., 2017; He et al., 2019; Hipp et al., 2012). Therefore, at sufficiently large distances where the percentage of zero- and near-zero phase-delay connectivity does not vary as a function of Euclidean distance, artefactual connectivity due to signal leakage is likely to be negligible. We examined this relationship graphically (Fig. 1a), finding that the proportion of zero- and near-zero phase-delay connectivity did not vary as a function of Euclidean distance > 60mm. As expected (Srinivasan et al., 2007), this was true for all frequency bands. This finding is consistent with artefactual connectivity due to signal leakage being likely negligible beyond 60mm, in keeping with previous estimates using simulated data (Anzolin et al., 2019; Gohel et al., 2017).

When limiting our analyses to region-pairs 80mm (where zero- / near-zero phase-delay connectivity was minimal) or more apart, we found that most functional connectivity still occurred with zero- and near-zero phase-delay, in all frequency bands except alpha (where 47% of all functional connections occurred with zero- or near-zero phase-delay, Fig. 1b).

Finally, we looked at homotopic interhemispheric functional connectivity as an example of connectivity known to robustly occur across species and measurement modalities (Cauda et al., 2021; Conturo et al., 1999; Engel et al., 1991; O’Reilly & Elsabbagh, 2021). Remarkably, 83-89% of homotopic interhemispheric connectivity occurred with zero- or near-zero phase-delays (Fig. 1b), despite the significant Euclidean distance between them. This proportion was greater than that of 34 randomly selected pairs of heterotopic interhemispheric region-pairs, in each frequency band (e.g. alpha: *X*^2^ (1, n = 5,202) = 8.19, *p* = .0042, Supplementary Table S1).

### Comparing the performance of zero-phase inclusive versus exclusive functional connectivity metrics

#### Test-retest reliability is higher in zero-phase inclusive than exclusive connectivity methods

In the alpha band, median edgewise ICC values were 0.20 – 0.71 across zero-phase-inclusive methods and 0.07 – 0.17 across zero-phase-exclusive methods (n = 99, Fig. 2, Supplementary Table S2). Absolute agreement was greatest in high-beta for all zero-phase-inclusive methods and Img COH. It was highest in alpha for the wPLI and in low-beta for Orth AEC. The same patterns were also found for the consistency of each measure (Supplementary Table S3).

**Fig. 2:**
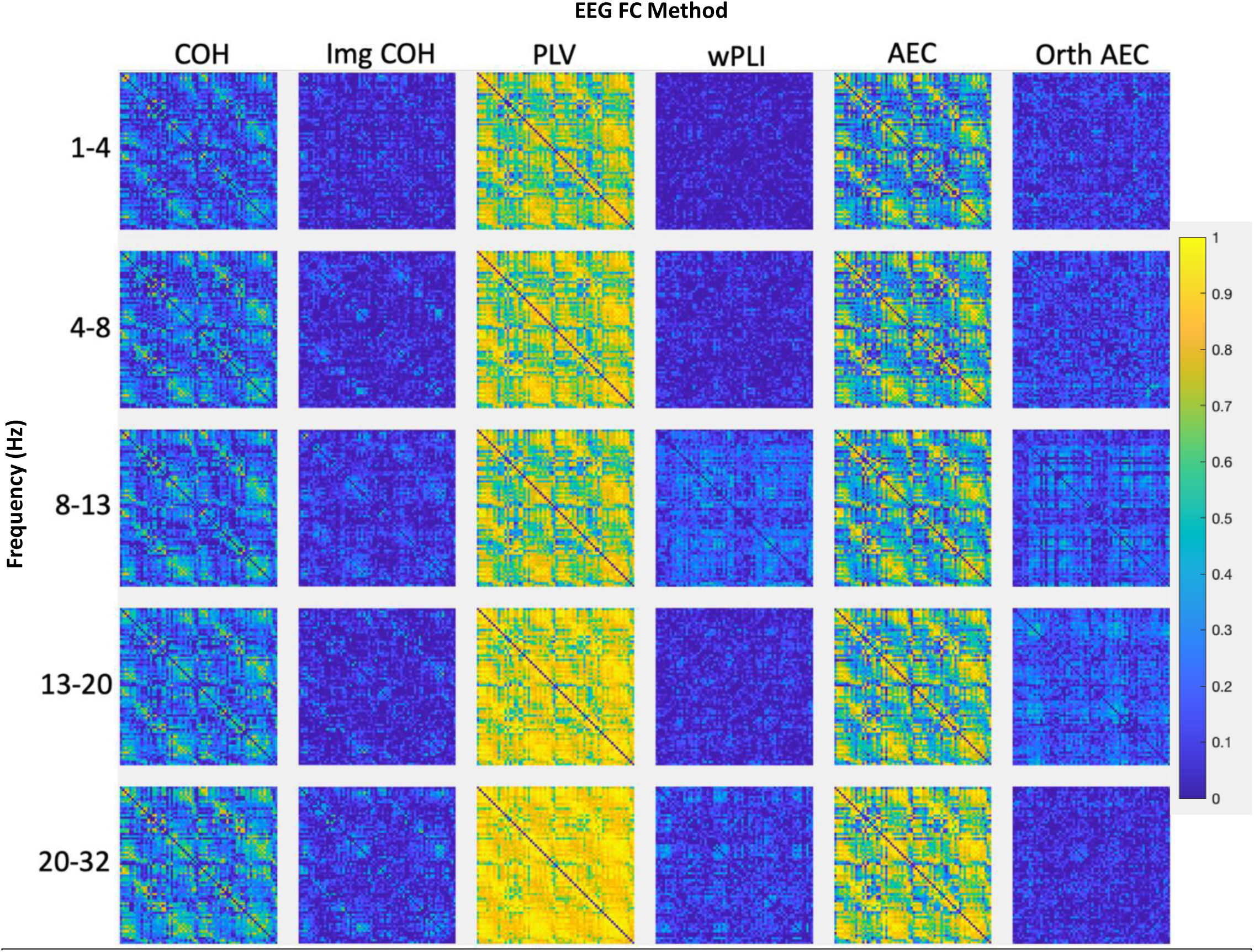
edgewise absolute agreement across functional connectivity methods and frequency bands. Edgewise absolute agreement, quantified by intraclass correlation coefficient (ICC), for each functional connectivity method. In 8 - 13 Hz, for example, median ICC values were 0.20 – 0.71 across zero-phase-inclusive methods versus 0.07 – 0.17 across zero-phase-exclusive methods. High absolute agreement was seen between homotopic interhemispheric region-pairs (off-diagonal edges in the adjacency matrix) in zero-phase-inclusive but not exclusive methods. High absolute agreement was seen between adjacent regions (main diagonal) in zero-phase inclusive methods, but not in zero-phase exclusive methods. Brain regions in the matrices were ordered by lobe and their proximity to each other. The colour bar denotes the edgewise ICC value. Negative ICC values were limited at 0.

We found high absolute agreement for measured connectivity between region-pairs with a small Euclidean distance between them using zero-phase inclusive but not exclusive methods (Fig. 2). There are two plausible causes for this: 1) neighbouring region-pairs reliably (structurally and functionally) connect to each other (Bullmore & Sporns, 2012; Conturo et al., 1999; Finger et al., 2016), and this functional connectivity occurs with zero- and near-zero phase-delays and 2) artefactual connectivity (Hipp et al., 2012).

Absolute agreement at homotopic interhemispheric region-pairs (Fig. 2) was high in all zero-phase inclusive methods, moderate with Img COH, and low with the Orth AEC and wPLI.

The relative test-retest reliability quantities across methods (Colclough et al., 2016; Fraschini et al., 2019) and frequency bands (van der Velde et al., 2019) is consistent with previous work, despite methodological differences.

#### Convergence with underlying neurobiology

We assessed the extent to which metrics derived from each functional connectivity method converged with underlying neurobiology by quantifying: 1) structure-function concordance, as functional connectivity is influenced by monosynaptic structural cortico-cortical pathways (Baum et al., 2020; Chu et al., 2015; Finger et al., 2016; Fotiadis et al., 2024; Honey et al., 2009), 2) the ability of functional connectivity metrics to capture homotopic interhemispheric connectivity, 3) their ability to predict participant age.

#### Concordance with structural connectivity is greater in zero-phase-inclusive methods

For all zero-phase-delay inclusive methods, mean structure-function concordance was significantly greater than zero in every frequency band (n = 50, one-sample t-test or Wilcoxon signed-rank test, *μ* = 0, *p* <.0001, Bonferroni corrected), Fig. 3. Concordance was not significantly different to zero for wPLI and orthogonalized AEC (one-sample t-test, *p* > .05). Surprisingly, concordance was significantly less than zero for Img COH (one-sample t-test or Wilcoxon signed-rank test, *p* <.0001, Bonferroni corrected), Fig. 3a-b. Concordance for each zero-phase-delay inclusive method was significantly higher than that for its methodologically corresponding zero-phase-delay exclusive method (paired-samples t-test or Wilcoxon signed-rank test, *p* < .0001, Bonferroni corrected, Fig. 3a-b). All findings were replicated when analysing each of the two sites separately, despite a difference in mean age between sites and large site effects in MRI analyses (Jovicich et al., 2006), Supplementary Table S4a-b.

**Fig. 3:**
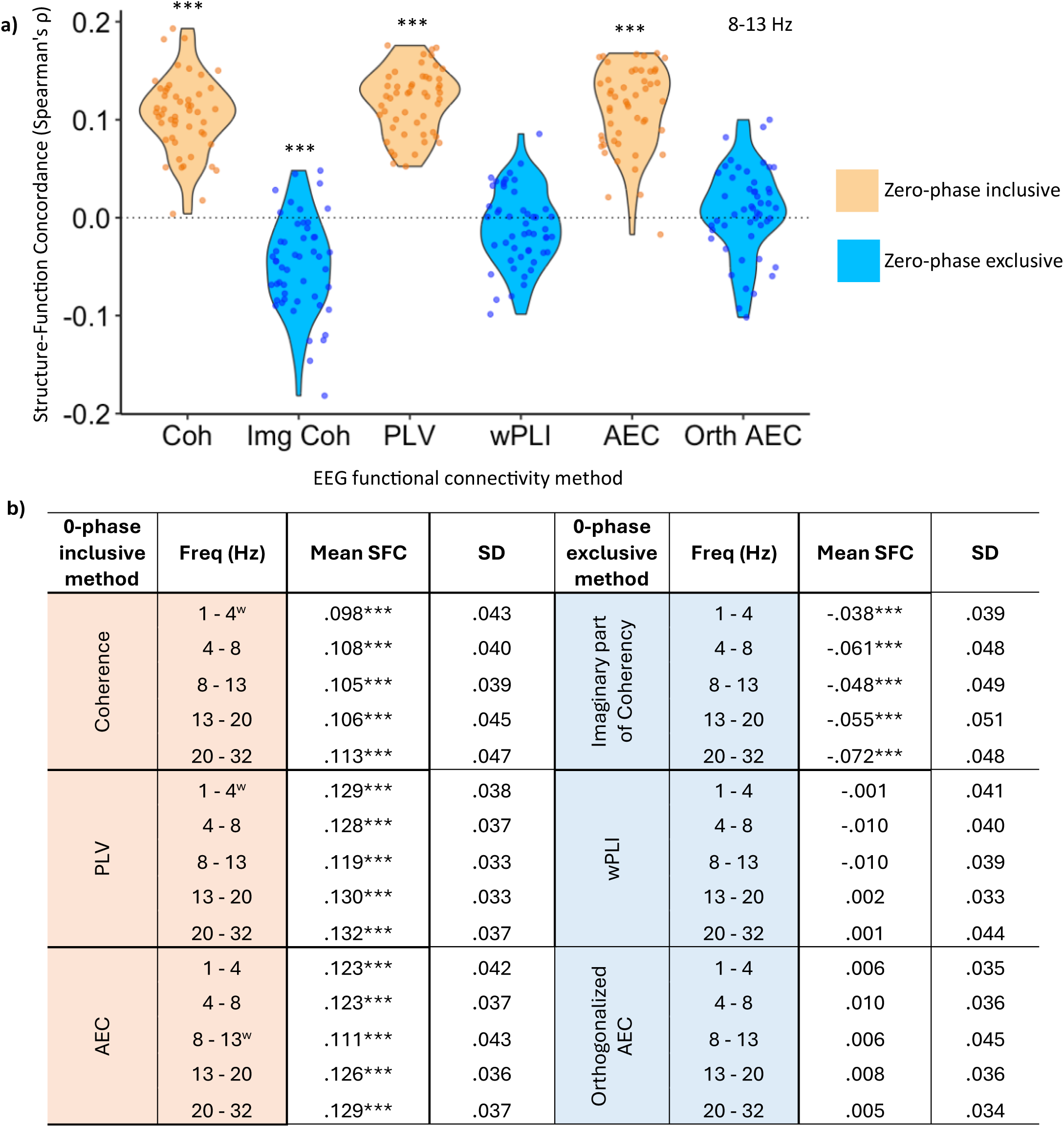
Structure-function concordance is higher in zero-phase inclusive versus exclusive methods. **a)** Structure-function concordance in the alpha band and **b)** in all frequency bands (n = 50). In all frequency bands, concordance was significantly greater than zero with zero-phase-inclusive methods (one-sided t-test or Wilcoxon sum-ranked test, *μ* = 0, *p* < .0001, Bonferroni corrected). Concordance was not significantly different from zero for two zero-phase-exclusive methods (wPLI and orthogonalized AEC). Surprisingly, concordance was significantly less than zero for the imaginary part of coherency. Concordance for each zero-phase-inclusive method was significantly higher than that of its methodologically corresponding zero-phase-exclusive method in each frequency band (paired samples t-test or Wilcoxon signed-rank test, *p* < .0001, Bonferroni corrected). *** concordance significantly different from zero, *p* < .0001, Bonferroni corrected. ^w^ = data not normally distributed, therefore one-sided Wilcoxon sign-rank test performed, otherwise one-sided t-tests performed. SFC = structure-function concordance.

#### Homotopic interhemispheric functional connectivity is robustly present with zero-phase-inclusive but not zero-phase-exclusive methods

Homotopic interhemispheric connectivity is robustly found across species and modalities (Engel et al., 1991; Mancuso et al., 2019; O’Reilly & Elsabbagh, 2021; Schmahmann & Pandya, 2009; Vicente et al., 2008; Witham et al., 2007). We examined each functional and structural connectivity method’s ability to capture this fundamental brain property. We calculated the edgewise mean functional (n = 153) and structural (n = 50) connectivity strength across all participants. On visual inspection, strong homotopic interhemispheric connectivity was found with all zero-phase-inclusive functional connectivity methods (Fig. 4a) and with structural connectivity (reflecting callosal connections, Fig. 4b). No zero-phase-exclusive methods showed prominent homotopic interhemispheric connectivity. Img COH showed significantly weaker connectivity between homotopic interhemispheric region-pairs than between non-homotopic region-pairs (Fig 4a, Supplementary Table S5), which we explored in a *post hoc* analysis below. Of note, given the relatively large Euclidean distances between them (median 57mm, IQR 21-80mm), functional connectivity between homotopic region-pairs would be minimally impacted by signal leakage.

**Fig. 4:**
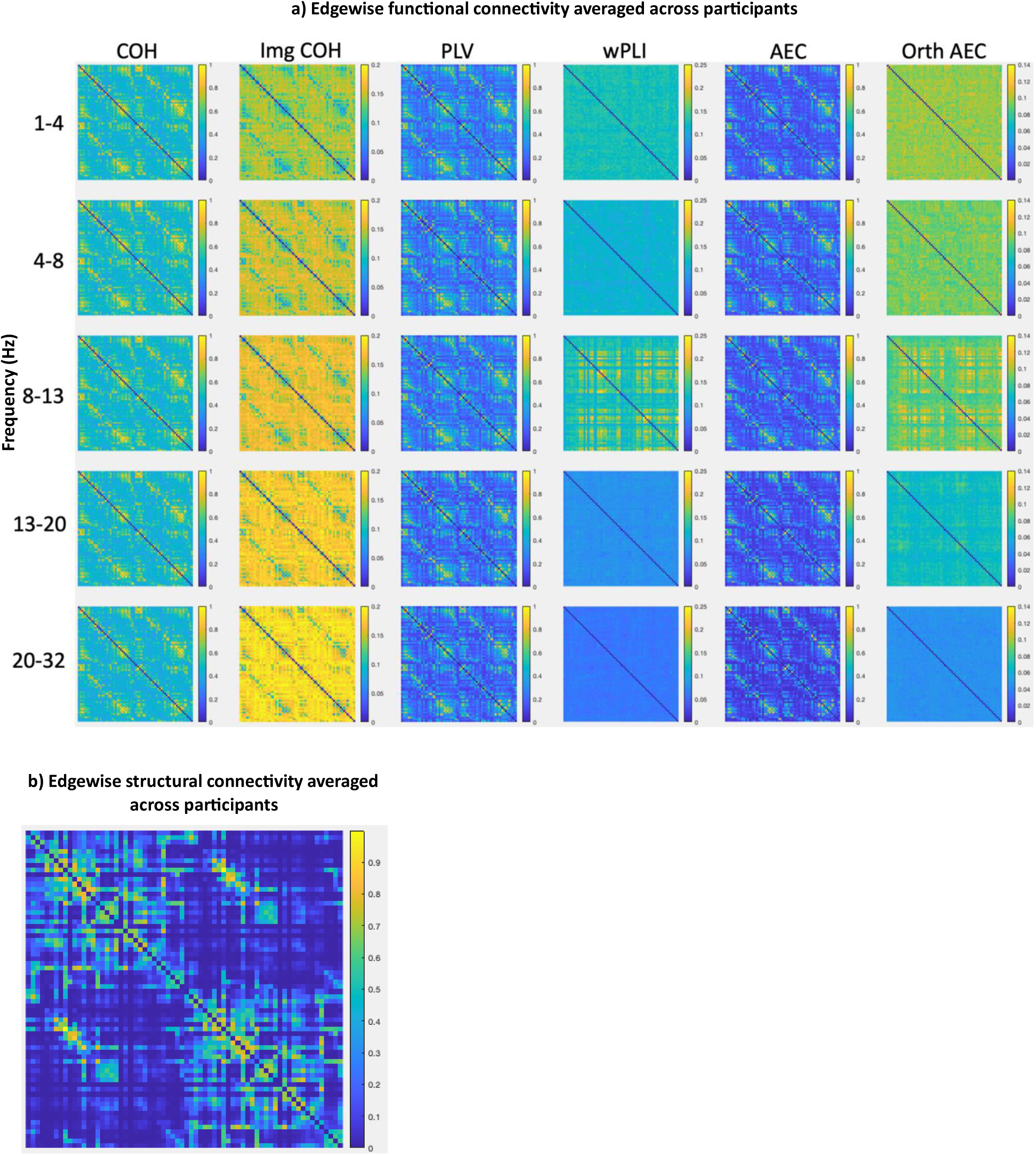
Edgewise structural and functional connectivity strengths. Mean edgewise **a)** EEG functional connectivity matrices (n = 153) and **b)** structural connectivity matrix (n = 50). Strong homotopic interhemispheric connectivity (seen at the off-diagonal edges in each adjacency matrix) was found with structural connectivity and with all zero-phase-inclusive functional connectivity methods. No zero-phase-exclusive methods showed prominent callosal connectivity. The imaginary part of coherency showed significantly weaker connectivity between homotopic interhemispheric region-pairs than between other region-pairs. Zero-phase-delay exclusive methods did not show strong connectivity between neighbouring region-pairs (main diagonal). Colour bars reflect connectivity strength, kept constant within each connectivity method; yellower colours denote higher connectivity strength. Brain regions in matrices are ordered by lobe and their proximity to each other.

#### The ability to predict current age is better in zero-phase-inclusive versus exclusive methods

Given that brain functional connectivity matures with age (Collin & Van Den Heuvel, 2013; Edde et al., 2021; Fair et al., 2009), we quantified the extent to which age-related changes are captured by each functional connectivity method. We assessed the ability of mean strength derived from each method to predict current age over and above head circumference (which correlates with age). First, we calculated the R^2^ from 10-fold cross-validation using head circumference as the only predictor of current age (model 1, cross-validation R^2^ = .27). Then, we added mean strength to this model (model 2). The subsequent change in R^2^ (model 2 – model 1) is reported in Table 3a, n = 145. Across all frequency bands, model 2 – model 1 R^2^ was greater in a zero-phase-inclusive method compared to its methodologically corresponding zero-phase-exclusive method; permutation analysis showed that this was statistically significant (false discovery rate corrected) in 10/15 comparisons.

**Table 3a:** Prediction of current age is greater in zero-phase inclusive versus exclusive methods. Values show model 2 – model 1 ten-fold cross validation R^2^ (n = 145) for each functional connectivity method and frequency band. Model 1: *current age ∼ head circumference* (model 1 cross-validation R^2^ = .27). Model 2: *current age ∼ head circumference + mean strength*.

| Freq (Hz) | 0-phase-inclusive method | Model 2 – 1 cross validation $R^2$ | 0-phase-exclusive method | Model 2 – 1 cross validation $R^2$ | $p_{FDR}$ permute |
| --- | --- | --- | --- | --- | --- |
| 1 - 4 | Coherence | .04 | Imaginary part of Coherency | .02 | .746 |
| 4 - 8 |  | .09 |  | .02 | **.041** |
| 8 - 13 |  | .06 |  | .05 | .905 |
| 13 - 20 |  | .07 |  | -.02 | **.020** |
| 20 - 32 |  | .01 |  | -.01 | .070 |
| 1 - 4 | PLV | .05 | wPLI | -.01 | **.010** |
| 4 - 8 |  | .12 |  | .02 | **.015** |
| 8 - 13 |  | .16 |  | .11 | .101 |
| 13 - 20 |  | .11 |  | .01 | **.041** |
| 20 - 32 |  | .10 |  | .04 | **.015** |
| 1 - 4 | AEC | .04 | Orthogonalized AEC | .00 | .285 |
| 4 - 8 |  | .10 |  | -.01 | **.041** |
| 8 - 13 |  | .15 |  | .02 | **.000** |
| 13 - 20 |  | .14 |  | .02 | **.041** |
| 20 - 32 |  | .15 |  | -.01 | **.000** |

#### Prognostic ability for longitudinal changes in spatial working memory is higher in zero-phase inclusive versus exclusive methods

We explored the ability of metrics derived from each functional connectivity method to act as prognostic biomarkers (Pletcher & Pignone, 2011) for longitudinal changes in spatial working memory ability, above and beyond current working memory abilities. We chose spatial working memory as an outcome measure because it has highly specific associations with functional connectivity (Dai et al., 2017b; Fox et al., 2005; Hampson et al., 2006; Kang et al., 2011) and altered working memory is implicated in several neuropsychiatric conditions (Alloway et al., 2009; Chai et al., 2018; Gathercole & Alloway, 2006).

We found significant pathlength x age interactions in general linear models predicting changes in spatial working memory performance, when analysing all 6-31-year-old participants together (examples for coherence and Img COH provided in Supplementary Tables S6-7). Therefore, children (6-12 years, n = 32), adolescents (13-17 years, n = 41) and adults (18-31 years, n = 36) were analysed separately. Model covariates (selected using Akaike information criterion) were kept constant within each age group.

A main effect of network pathlength was only found for methods sensitive to zero-phase-delay connectivity: coherence in children (β = 3.8, SE = 1.5, *p* = .014; full model F_3/28_ = 4.01, *p* = .017) and PLV in adults (β = −2.6, SE = 1.1, *p* = .027; full model F_3/32_ = 18.53, *p* = .003), Supplementary Tables S8-9. For these statistically significant findings, we used leave-one-out cross-validation (LOOCV) to quantify model performance (Table 3b). First, we calculated the LOOCV R^2^ using covariates (model 1). Then, we added weighted, normalised pathlength to this model (model 2). Model 2 – model 1 LOOCV R^2^ increased by .101 for coherence in children and .074 for PLV in adults. For comparison in their methodologically corresponding zero-phase-exclusive methods, model 2 – model 1 LOOCV R^2^ *decreased* by .005 for Img COH and .036 for wPLI.

**Table 3b:** Prediction of longitudinal changes in spatial working memory performance is greater in zero-phase inclusive versus exclusive methods. In models predicting longitudinal changes in spatial working memory performance, a significant main effect of functional connectivity derived pathlength was only found for coherence in the 6–12-year group and PLV in the 18-31-year group. For these models, and their corresponding zero-phase-exclusive methods, we report model 2 – model 1 leave-one-out cross-validation R^2^. Model 1: *longitudinal change in spatial working memory ∼ covariates*. Model 2: *longitudinal change in spatial working memory ∼ covariates + functional connectivity derived pathlength*. *significant main effect of covariate in general linear model (*p* < .05). SWM = spatial working memory, T1 = timepoint 1, T2 = timepoint 2, PL = pathlength

| Age Group | Model 1 covariates | Model 1 $R^2$ | Covariate added in model 2: | Model 2 $R^2$ | Model 2 – Model 1 $R^2$ |
| --- | --- | --- | --- | --- | --- |
| 6-12 yrs<br>($n = 32$) | Age + T1 SWM | .006 | Coh PL* | .107 | .101 |
|  |  |  | Img COH PL | .001 | -.005 |
| 18-31 yrs<br>($n=36$) | Age + T1 SWM + T2-T1 time interval | .489 | PLV PL* | .563 | .074 |
|  |  |  | wPLI PL | .453 | -.036 |

### *Post hoc* analyses

An unexpected finding from our analyses was negative structure-function concordance with Img COH. We theorised that this was because the strongest structural connections had the most ‘penalised’ functional connections when using Img COH, based on the following logic: stronger structural connections result in quicker (decreased) signal transmission time between region-pairs (Westerhausen et al., 2006; Whitford et al., 2011). Quicker transmission time means that oscillatory activity at each connected region-pair occurs with smaller time-delays, and therefore, smaller phase-delays. Zero-phase-exclusive methods may penalize, rather than merely exclude, functional connectivity at near-zero-phase-delays (a salient methodological detail is that zero-phase-exclusive methods modify the connectivity strength value based on the magnitude of phase delay, while zero-phase-inclusive methods to do not). Hence, stronger structural connections may lead to weaker functional connections with some zero-phase-exclusive methods.

To test this, we hypothesised that: 1) the connectivity strength values of zero- and near-zero phase-delay connections are penalised, and not merely excluded, if derived from zero-phase-exclusive methods. 2) Stronger structural connections lead to functional connections with more penalizable phase delays, mediated by quicker possible white matter transmission time. We expected a partial mediation, as strong structural connections such as the corpus callosum also produce zero-phase-delay functional connectivity through resonance-induced synchrony. 3) Structural connections that allow the quickest connectivity between region-pairs result in the weakest functional connectivity if derived from Img COH.

#### Near-zero-phase-delay functional connectivity may be penalised and not merely excluded by zero-phase-delay-exclusive methods

We inspected the relationship between absolute phase delay and functional connectivity strength (n = 153), between region-pairs where signal leakage is likely negligible (region-pairs ≥80mm apart), Fig. 5. With zero-phase-inclusive methods, the median connectivity strength was significantly higher for near-zero/near-π phase-delay connectivity compared to connectivity at other phase delays, in all frequency bands (Fig. 5b). For Img COH and wPLI the opposite was true: median connectivity strength was significantly lower for near-zero/near-π phase-delay connectivity compared to connectivity at other phase delays. For orthogonalized AEC, median connectivity strength was similar between near-zero phase-delay and other phase-delay connectivity, in keeping with the milder form of orthogonalization we used compared to others (Brookes et al., 2012; Hipp et al., 2012). Therefore, zero- and near-zero phase-delay functional connectivity may be penalised, and not merely excluded, by zero-phase-exclusive methods.

**Fig. 5:**
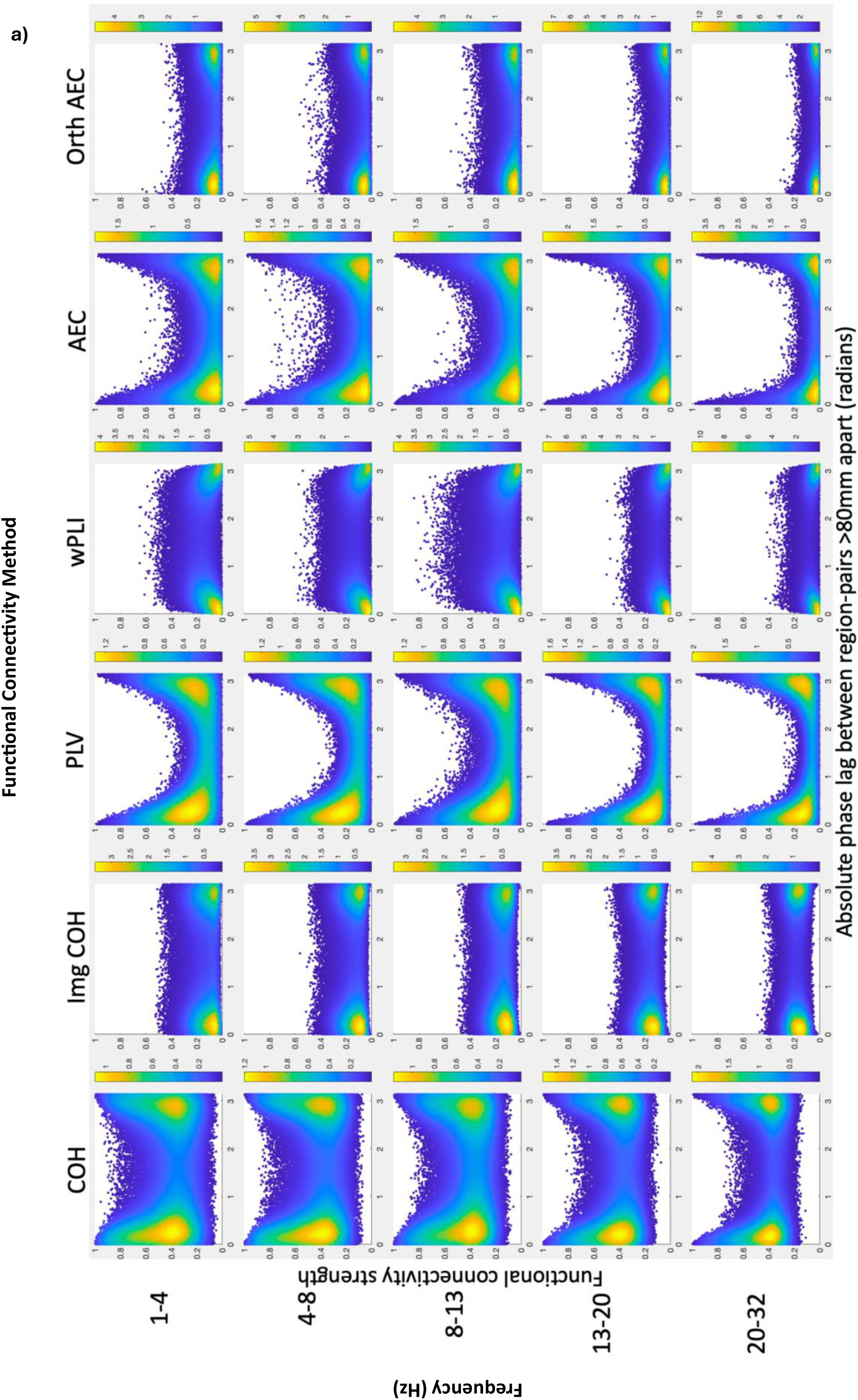
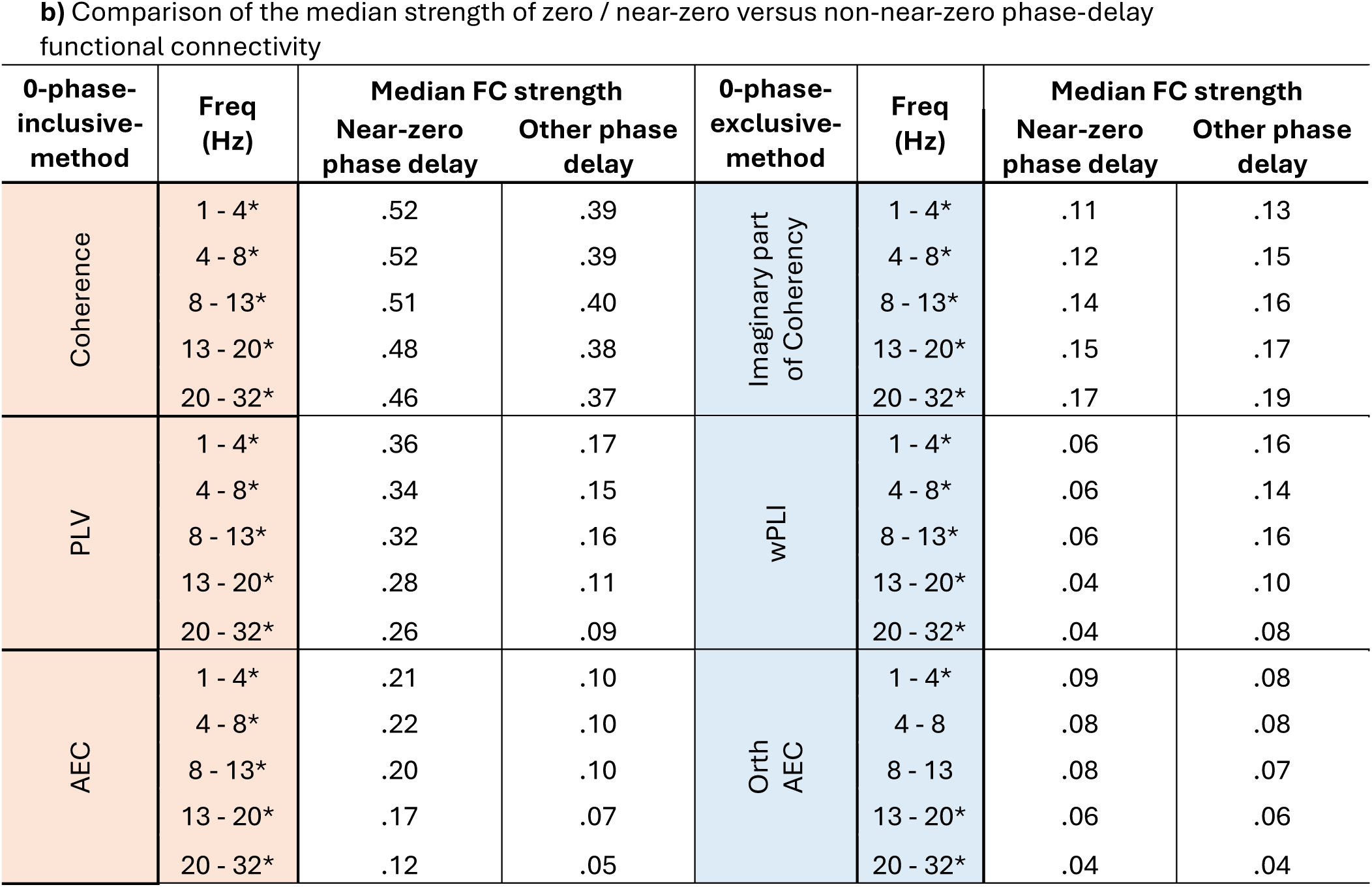
When signal leakage is likely negligible, most functional connectivity occurs with near-zero-phase-delay, penalised by zero-phase-exclusive methods. **a)** The relationship between absolute phase-delay and functional connectivity strength (yellower colours denote higher density of scatter points), and **b)** a comparison of edge strength when functional connectivity has zero / near-zero versus non-near-zero phase delays, for region-pairs > 80mm Euclidean distance apart (where artefactual connectivity due to signal leakage is likely negligible). For zero-phase-inclusive methods, across all frequency bands, median strength was significantly higher for near-zero-phase-delay connections compared to connections with non-near-zero-phase-delays. For the imaginary part of coherency and wPLI, across all frequency bands, the median strength was significantly lower for near-zero phase-delay connectivity compared to non-near-zero-phase-delay connectivity. For orthogonalized AEC, median strength was similar between near-zero and non-near-zero phase delay connections. Zero-phase-exclusive methods modify the connectivity strength value based on the magnitude of phase delay, while zero-phase-inclusive methods to do not. Therefore, zero-phase-exclusive methods may penalise, rather than merely exclude, zero- and near-zero phase-delay connections. * *p* < .0001, Bonferroni corrected. FC = functional connectivity.

#### Stronger structural connectivity leads to functional connectivity with more penalizable phase-delays, mediated by shorter white matter transmission times

We hypothesised that the stronger a structural connection, the more likely it was to produce functional connectivity with phase-delays closer to zero, penalised by zero-phase-exclusive methods. The mean correlation between streamline count and penalisability of phase-delays was significantly greater than 0 for all frequency bands (n = 50, one-sample t-test, *μ* = 0, *p* < .0001, Bonferroni corrected), Fig. 6a. Hence, stronger structural connectivity led to functional connectivity with phase-delays more penalizable by zero-phase-exclusive methods.

**Fig. 6:**
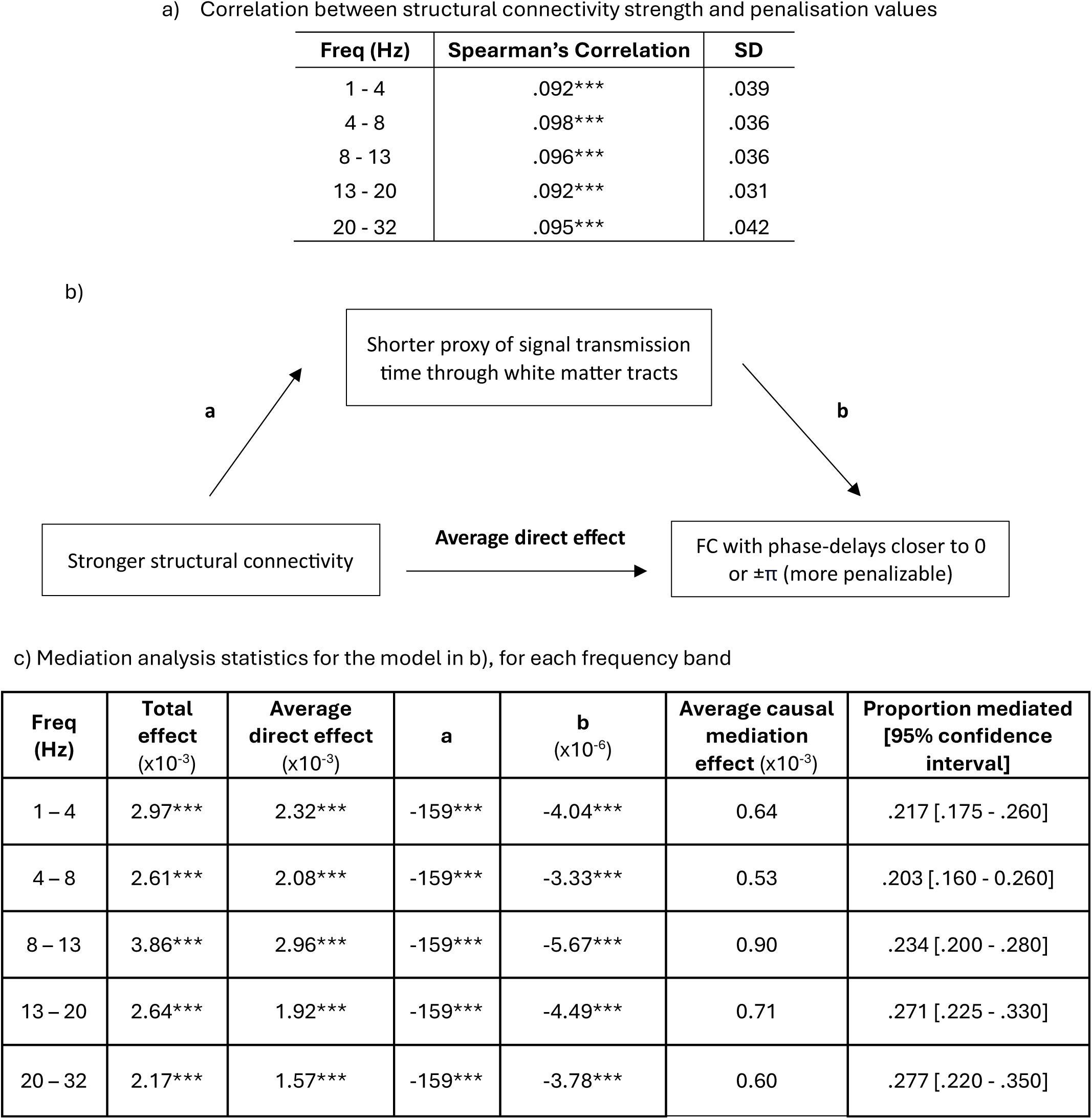
Stronger structural connectivity leads to functional connectivity with phase-delays closer to 0 or ±π, mediated by a proxy of shorter white matter transmission time. **a)** We created a penalisation scale between 0 and 1, where phase-delays most penalised by zero-phase-exclusive methods (0 and ±π) have a value of 1 and phase-delays least penalised (±π/2) have a value of 0. The mean correlation across participants between streamline count and penalisability of phase-delays was significantly greater than 0 for all frequency bands, n = 50 (*** *p* <.0001, Bonferroni corrected, one-sample t-test, *μ* = 0). **b-c)** We created a proxy of signal transmission time through white matter tracts. The effect of structural connectivity strength on the penalisabiltiy of the phase-delay of functional connectivity was partially mediated by the proxy of white matter transmission time, across all frequency bands. All effects were statistically significant (*** = *p* < .0001, Bonferroni corrected). SD = standard deviation.

To test whether this relationship was mediated by quicker signal transmission time through white matter connections, we used a proxy of signal transmission time through white matter tracts. Across all frequency bands, we found that the effect of structural connectivity strength on the penalisability of the phase-delay of functional connectivity was significantly partially mediated by the proxy of white matter transmission time (e.g. proportion of the direct effect mediated by the proxy of signal transmission time was .234 [95% confidence intervals, calculated using 1000 bootstrapped samples = .197 - .270] in alpha, Fig. 6b-c). The mediation was strongest in higher frequency bands, consistent with stronger structure-function concordance in higher frequency bands in zero-phase-inclusive methods (Fig. 3b).

#### Structural connections with shorter signal transmission time are associated with weaker functional connectivity with some zero-phase-exclusive methods

We found that the shorter the proxy of signal transmission time through white matter tracts, the stronger the functional connectivity value was for all zero-phase-inclusive methods (one-sample t-test, *μ* = 0, *p* < .0001, Bonferroni corrected), Fig. 7. This is the expected biological relationship (Bassett & Bullmore, 2017; Bullmore & Sporns, 2012; Cabral et al., 2014). This was also the case for the alpha band in wPLI (*p* < .05, Bonferroni corrected) and orthogonalized AEC (*p* < .001, Bonferroni corrected), but no other frequency bands when using these methods. Note that alpha has the smallest proportion of near-zero-phase-delay connectivity (Fig. 1b). For Img COH, the shorter the proxy of signal transmission time through white matter tracts, the weaker functional connectivity strength, in all frequency bands (one-sample t-test, *μ* = 0, *p* < .0001, Bonferroni corrected). This is inconsistent with prior network literature (Bassett & Bullmore, 2017; Bullmore & Sporns, 2012; Cabral et al., 2014).

Thus, the structural connections that are prioritised in brain through quick structural pathways may be penalised by some zero-phase-delay-exclusive functional connectivity methods.

**Fig. 7:**
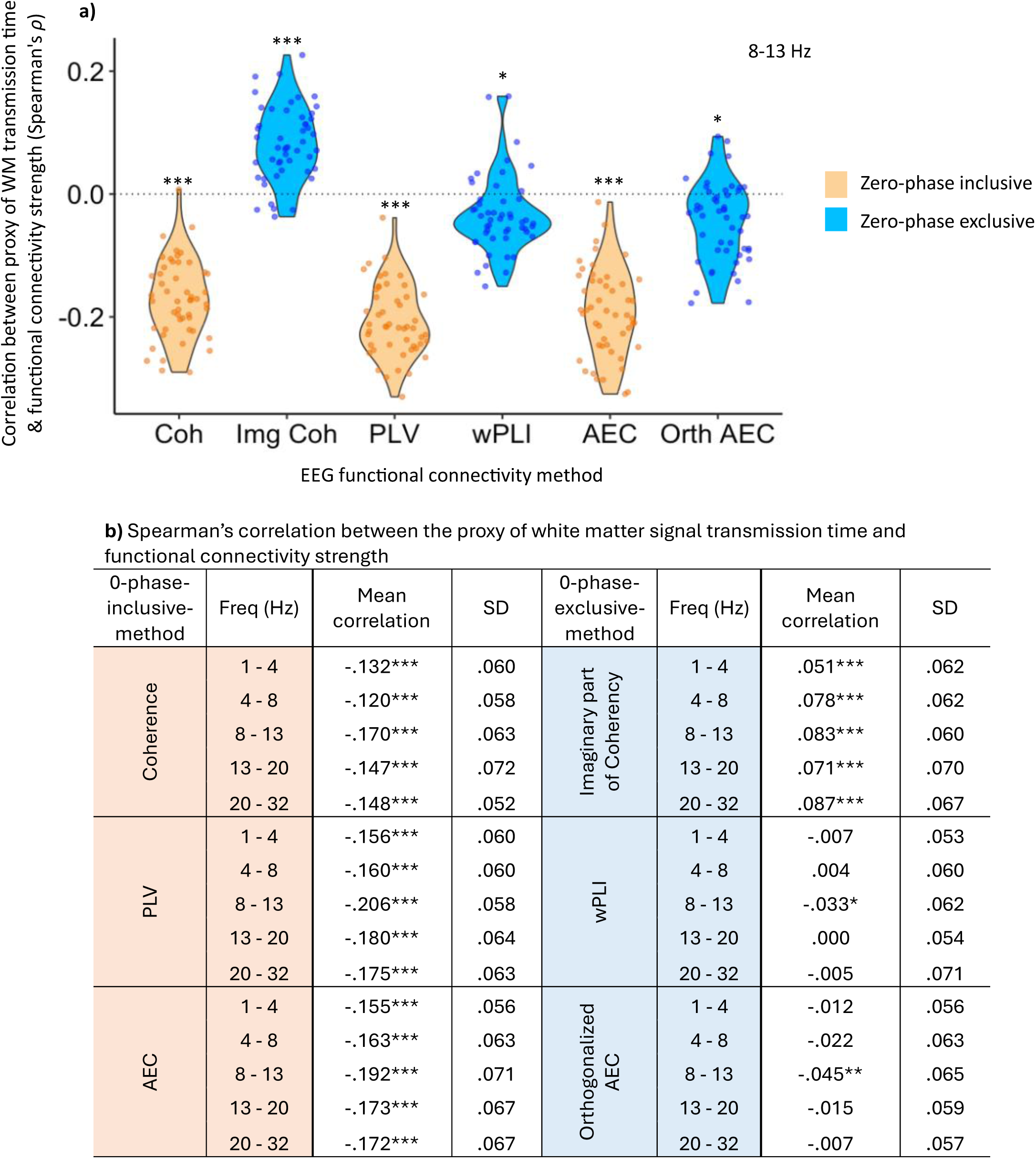
Structural connections with shorter signal transmission time are associated with weaker functional connectivity with some zero-phase-exclusive methods. We created a proxy of signal transmission time through white matter tracts. The quicker (shorter) the proxy of signal transmission time through structural connections, the stronger the functional connectivity value for all zero-phase-inclusive methods (one-sample t test, *μ* = 0, *p* < .0001, Bonferroni corrected), **a)** shown graphically for alpha and **b)** tabulated for all frequency bands (n = 50). This was also the case for alpha (but not other frequency bands) in wPLI (*p* < .05, Bonferroni corrected) and orthogonalized AEC (*p* < .001, Bonferroni corrected). For the Img COH, the quicker the proxy of structural transmission time, the weaker functional connectivity strength, in all frequency bands (one-sample t-test, *μ* = 0, *p* < .0001, Bonferroni corrected). Such a relationship is inconsistent with prior literature. \*\*\**p* < .0001, \*\**p* < .001, \**p* < .05 (Bonferroni corrected).

## Discussion

We demonstrated that most interactions between cortical region-pairs occur with zero- and near-zero phase-delay. We used a novel approach to show that this is also the case when artefactual connectivity due to signal leakage is likely negligible. Including zero- and near-zero phase-delay connectivity increased test-retest reliability, convergence with neurobiology (concordance with structural connectivity, ability to capture homotopic interhemispheric connectivity and ability to predict chronological age) and prognostic ability for longitudinal changes in cognitive ability. Thereby, through converging lines of evidence, we show that functional connectivity metrics that include zero- and near-zero phase-delay connections have more properties desired in candidate biomarkers. We propose that the decreased structure-function concordance, ability to capture age, and ability to predict longitudinal changes in cognition associated with zero-phase-exclusive methods are due to their penalisation of most ‘true’ cortico-cortical functional connectivity.

We investigated homotopic interhemispheric connectivity as an example of structural and functional connectivity robustly found across species (Engel et al., 1991; Mancuso et al., 2019; O’Reilly & Elsabbagh, 2021; Schmahmann & Pandya, 2009; Vicente et al., 2008; Witham et al., 2007). We captured conspicuous homotopic connections using structural connectivity and zero-phase-inclusive functional connectivity methods. We found that they were either inconspicuous or penalised when using zero-phase-exclusive methods. In contrast, a previous study (Hipp et al., 2012) captured connectivity between sensory homotopic region-pairs using orthogonalized AEC. However, given our and others’ (Engel et al., 1991; O’Reilly & Elsabbagh, 2021) finding that homotopic region-pairs predominantly interact with zero- and near-zero-phase-delay, our findings are expected. We see the inability to capture this fundamental feature of brain connectivity as a key weakness of zero-phase-exclusive methods.

We found that structure-function concordance was significantly greater when using zero-phase-inclusive compared to zero-phase-exclusive methods. This is consistent with a previous small study of 17 participants (Finger et al., 2016). Additionally, we were surprised that with some zero-phase-exclusive methods, here Img COH, the strongest structural connections paradoxically had the weakest functional connections. We showed that this was because stronger structural connectivity led to functional connectivity with phase-delays closer to zero, mediated by a proxy of shorter signal transmission times. Then, we showed that functional connectivity at near-zero-phase-delays had penalised strength when derived from some zero-phase-exclusive methods. Hence, for Img COH, we found that region-pairs connected by white matter tracts with the quickest possible signal transmission times had the weakest functional connections. This relationship is inconsistent with previous network literature (Bassett & Bullmore, 2017; Bullmore & Sporns, 2012; Cabral et al., 2014).

Predictive validity for longitudinal changes in spatial working memory performance, as well as the ability to predict current age, was substantially greater when zero- and near-zero phase-delay connections were included. Strengths of our approach included the fact that our spatial working memory prediction findings were derived using region-pairs (Almeida et al., 2015; Jones & Wilson, 2005; MacKey & Curtis, 2017; Salimi et al., 2021; Siapas et al., 2005; Wager & Smith, 2003; Wirt & Hyman, 2017), the frequency band and the network topology measure known to be implicated in spatial working memory (Bauer et al., 2021b; Dai et al., 2017a; Fell & Axmacher, 2011; Hyman et al., 2010; Jones & Wilson, 2005; Siapas et al., 2005), specified *a priori*. Consistent with our findings, a small, cross-sectional prior study (Jian et al., 2017) found that including versus excluding zero-phase-delay connectivity substantially increased the variance explained in predicting current task state. It showed that predictive ability was not dependent on differing characteristics between functional connectivity methods besides the inclusion or exclusion of zero-phase-delay connectivity: it excluded zero-phase-delay connectivity using a large-Laplacian while keeping the connectivity method constant.

Conversely, zero-phase-exclusive methods did demonstrate some advantages. In agreement with others (Hipp et al., 2012; Nolte et al., 2004; Vinck et al., 2011), it appeared that zero-phase-exclusive methods likely minimised artefactual connectivity due to volume conduction and signal leakage: functional connectivity between neighbouring region-pairs was minimal when using them (Fig. 4). However, valid functional connections are also most commonly present between nearby region-pairs (Bassett & Bullmore, 2017; Honey et al., 2009; Mehrkanoon et al., 2014; Varela et al., 2001), and were not reliably captured by zero-phase-exclusive methods (Figs. 2 and 4).

Quantifying ‘true’ zero- and near-zero phase-delay connectivity has not been previously achieved in empirical EEG or MEG data. Our approach to identify the distance between region-pairs where the effects of signal leakage were likely negligible was novel. Previous approaches have used simulated data (Anzolin et al., 2019; Gohel et al., 2017), with findings consistent with ours. However, simulated data cannot yet accurately inform us about the proportion of ‘true’ cortico-cortical functional connectivity that occurs with zero- and near-zero phase-delay, as even the most biologically realistic simulations (Moran et al., 2013) cannot accurately model the emergent properties of whole-cortex connectivity (Engel et al., 1991; Mill et al., 2017; Uhlhaas et al., 2009). Therefore, unlike simulated data approaches, our approach can illustrate that most cortical functional connectivity occurs at zero- or near-zero phase-delay, and that excluding it has deleterious effects on the properties of biomarkers derived from functional connectivity metrics. While the extent of artefactual zero-phase-delay connectivity in empirical signals is dependent on methodological choices, our findings highlight that zero- and near-zero-phase delay synchronisation comprises a substantial proportion of ‘true’ neural interactions.

A key strength of our functional connectivity metric performance assessment approach was the use of convergence across performance criteria – each based on different modalities not trivially correlated with each other. Previous attempts at assessing the validity of zero-phase-inclusive versus zero-phase-exclusive methods have used single measures of functional connectivity performance (Brookes et al., 2012; Colclough et al., 2016; Hipp et al., 2012), which may lead to bias (Fraschini et al., 2019; Noble et al., 2021). Even using (only) two modalities could be misleading: for example, both structural connectivity and functional connectivity from zero-phase-inclusive methods are more reliable with decreasing Euclidean distance between region-pairs (Calamante, 2019; Jbabdi & Johansen-Berg, 2011). Thus, high structure-function concordance may reflect reliable but artefactual functional connectivity correlating with reliable and valid structural connectivity at small distances. Therefore, our use of non-neuroimaging markers of performance was important.

An additional strength was our multi-site study design; our most unexpected finding (negative structure-function concordance with Img COH) was reproduced when analysing each study site separately. Next, as a measure to assess convergence against, the structural connectome has high inter- and intra-individual reliability (Chu et al., 2015; Dennis et al., 2012). Further, to test each functional connectivity metric property, we used multiple analysis approaches, with consistent results.

Is high test-retest reliability a good measure of functional connectivity metric performance? High reliability has been used as evidence of the presence of volume conduction or signal leakage artefact in functional connectivity analyses (Colclough et al., 2016; Nagy et al., 2024). Further, high validity may be present with low test-retest reliability: brain-regions functionally connected at one moment may vary from the next. However, in the context of a candidate prognostic or diagnostic biomarker, high test-retest reliability is crucial; it establishes the upper limit of its predictive validity (Carmines & Zeller, 1979).

While we aimed to identify where the effects of signal leakage were negligible, the use of EEG additionally introduced volume conduction artefact (Wolters et al., 2004). In sensor space, artefactual connectivity due to volume conduction has a non-linear relationship with scalp distance (Srinivasan et al., 2007), making it challenging to account for. Source reconstruction aims to limit volume conduction artefact (He et al., 2019): in source space, simulated data show that artefactual connectivity due to volume conduction decreases with increasing Euclidean distance (Anzolin et al., 2019). This makes our approach to determine areas of minimal signal leakage also relevant to volume conduction. Further, EEG is a ubiquitous tool in research and clinical settings (Mclane et al., 2015), a strength when attempting to identify candidate biomarkers.

To some extent, our findings are surprising given the field’s move towards favouring zero-phase-exclusive methods (Garcés et al., 2022; Hipp et al., 2012; Miljevic et al., 2022; Nolte et al., 2004). Zero-phase-exclusive methods were introduced to identify ‘true’ brain interactions as opposed to trivial artefact (Nolte et al., 2004; Vinck et al., 2011). Over the last 20+ years, they have consistently been published as improvements on their corresponding zero-phase-inclusive methods. This position has been adopted by several studies attempting to identify candidate diagnostic or prognostic biomarkers (Garcés et al., 2022; Miljevic et al., 2022; Sato et al., 2023). Some authors have argued that zero-phase connectivity in the brain is physiologically implausible (Miljevic et al., 2022), despite robust evidence to the contrary.

The presence of ‘true’ zero-phase-delay synchrony in brain is unintuitive: action potential transmission delays due to axonal conduction and synaptic transmission are in the order of tens of milliseconds (Uhlhaas et al., 2009; Vicente et al., 2008). Such time-delays would cause substantial phase-delays between connected brain regions.

Thus, zero-phase-delay functional connectivity across distant neural populations would not be possible if the brain was a simple linear system. A leading postulated mechanism for zero-phase-delay connectivity is “resonance-induced synchrony” (Gollo et al., 2014): bidirectional information transfer between regions acts to mutually alter their dynamics into a stable zero-phase-delay pattern. Key structures have empirically shown to facilitate this, including cortico-thalamo-cortical loops (Uhlhaas et al., 2009; Vicente et al., 2008) and the corpus callosum (Bloom & Hynd, 2005; Musgrave & Gloor, 1980; Vicente et al., 2008; however see Contreras et al., 1996). Other hypothesised mechanisms of zero-phase-delay synchrony include entrainment (Gray & McCormick, 1996) and emergence (Kopell et al., 2000). It is possible that multiple mechanisms work in unison (Vicente et al., 2008).

The debate central to our study is the extent to which measured zero- and near-zero phase-delay connectivity in source reconstructed data is artefactual versus ‘true’. We found that most functional connectivity occurred with zero- and near-zero phase-delays, even when signal leakage was likely negligible. Converging lines of evidence demonstrate that including zero-phase-delay functional connectivity increases the performance of functional connectivity metrics as biomarkers. These findings challenge generally accepted assumptions that zero-phase-exclusive methods are superior to zero-phase-inclusive methods.

## Data and code availability

The LEAP Study data is available here, subject to an approved data application: https://redcap.pasteur.fr/surveys/?s=YRFF78PH89

Code will be made available here: https://github.com/cmehra-brain

## Author Contributions

Chirag Mehra – Conceptualisation, Methodology, Formal analysis, Writing - Original Draft, Writing - Review & Editing, Visualization

Ahmad Beyh - Formal analysis, Writing - Original Draft, Writing - Review & Editing, Visualization

Petroula Laiou – Resources, Writing - Original Draft

Pilar Garces - Data Curation, Formal analysis, Writing - Review & Editing

Emily JH Jones – Writing - Review & Editing, Funding acquisition, Project administration Luke Mason - Data Curation, Project administration

Jan Buitelaar – Writing - Review & Editing, Funding acquisition, Project administration Mark H Johnson - Project administration

Declan Murphy – Funding acquisition, Project administration

Eva Loth – Writing - Review & Editing, Funding acquisition, Project administration Flavio Dell’Acqua – Formal analysis, Writing - Review & Editing, Funding acquisition, Project administration, Supervision

Joshua B Ewen – Conceptualisation, Writing - Review & Editing, Supervision

Mark P Richardson - Writing - Review & Editing, Supervision, Funding acquisition Jonathan O’Muircheartaigh - Writing - Review & Editing, Supervision, Funding acquisition

## Acknowledgments

We are grateful to the AIMS-2-TRIALS LEAP-group for data collection and quality control procedures. We are grateful to Dr Piotr J. Franaszczuk for his expert advice on signal analysis and filtering and to Prof Dr Christine Ecker and team for pre-processing the T1-weighted MRI images.

## Supplementary Materials

**Supplementary Table S1.**
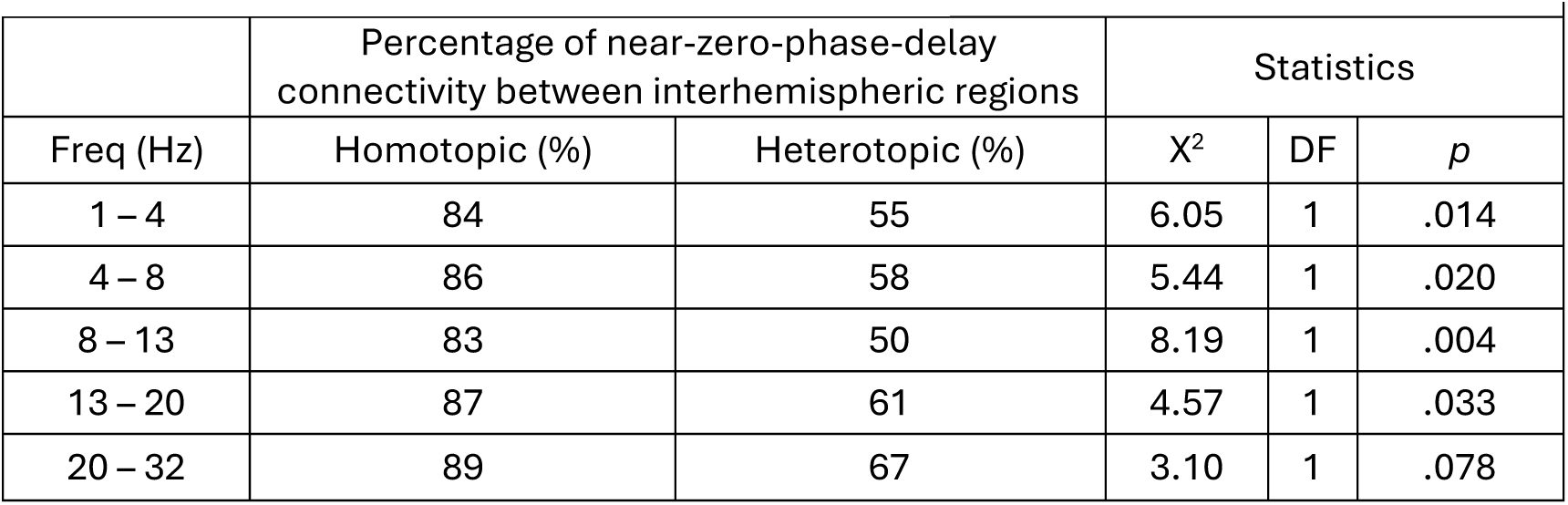
Comparing the percentage of near-zero-phase-delay functional connectivity between homotopic- and heterotopic-interhemispheric region-pairs. As there were 34 cortical regions per hemisphere, we randomly selected 34 heterotopic region-pairs, kept consistent across participants, to compare with the 34 homotopic region-pairs. DF = degrees of freedom.

**Supplementary Table S2.**
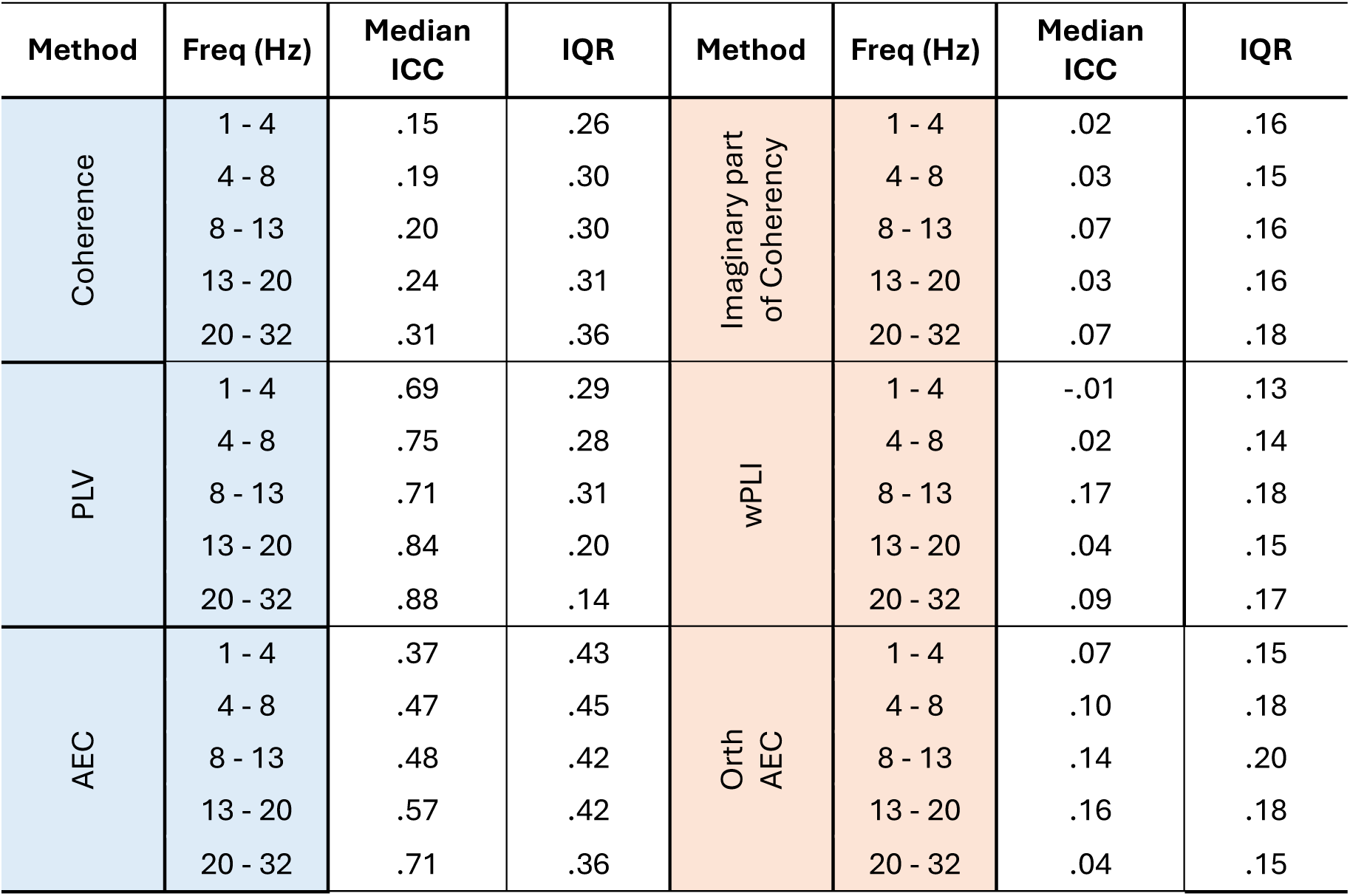
Intrasession edgewise absolute agreement of EEG functional connectivity methods. Median edgewise absolute agreement was calculated for each edge, using the interclass correlation coefficient (2,1; ICC). n = 99.

**Supplementary Table S3.**
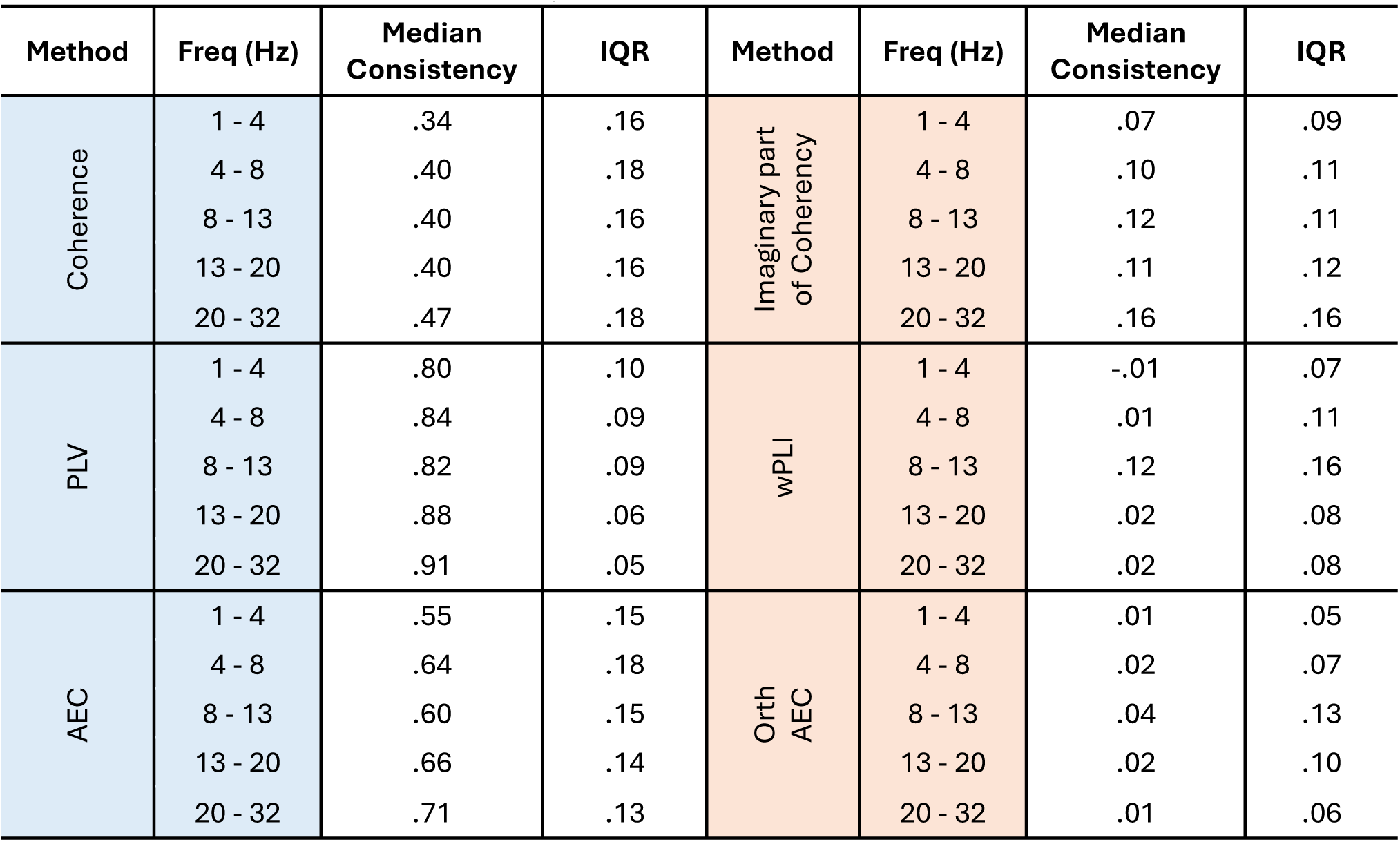
Intrasession consistency of the EEG functional connectivity adjacency matrix. Zero-phase-inclusive methods had moderate to high consistency values, while zero-phase-exclusive methods had near 0 consistency values.

**Supplementary Table S4.**
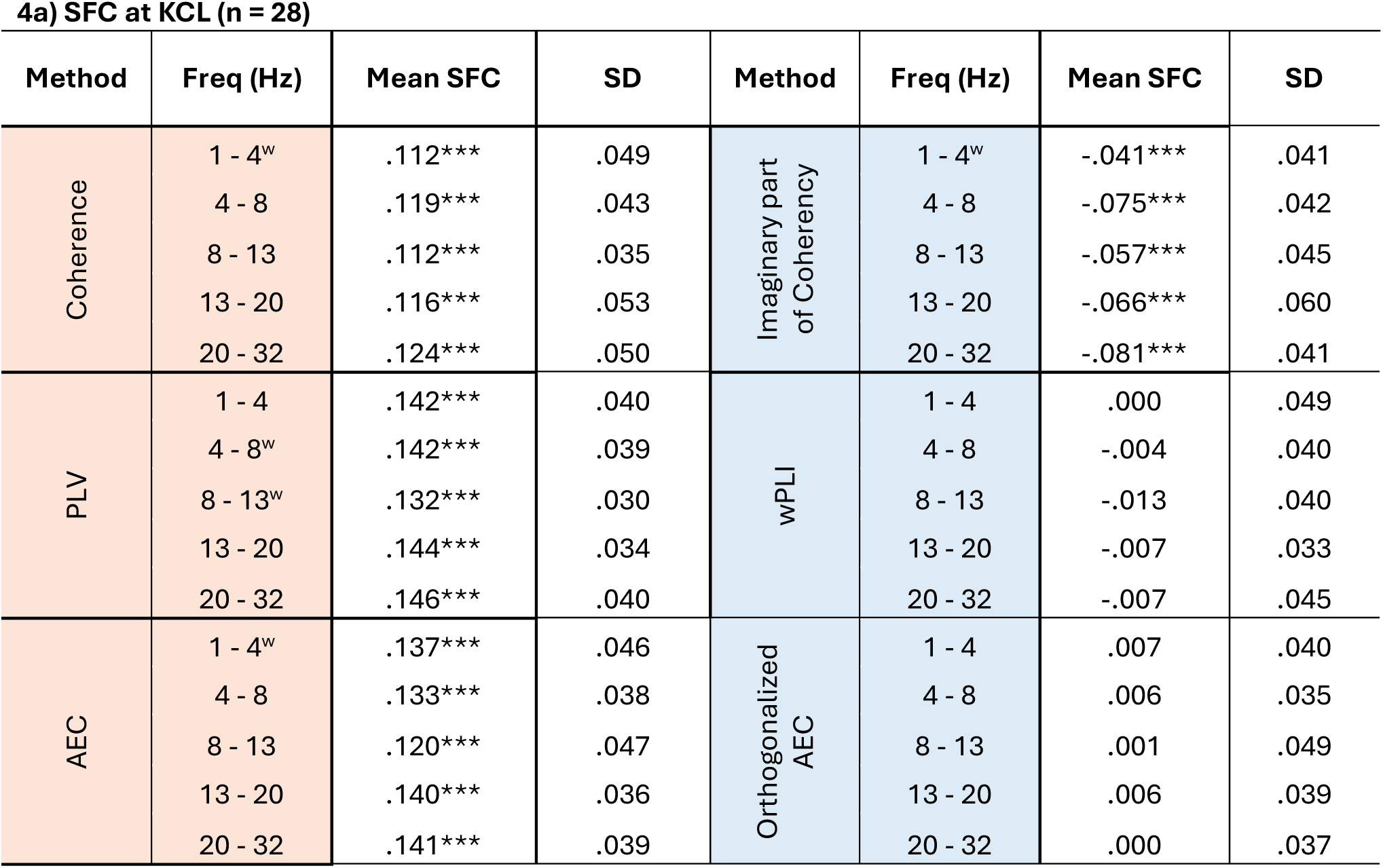
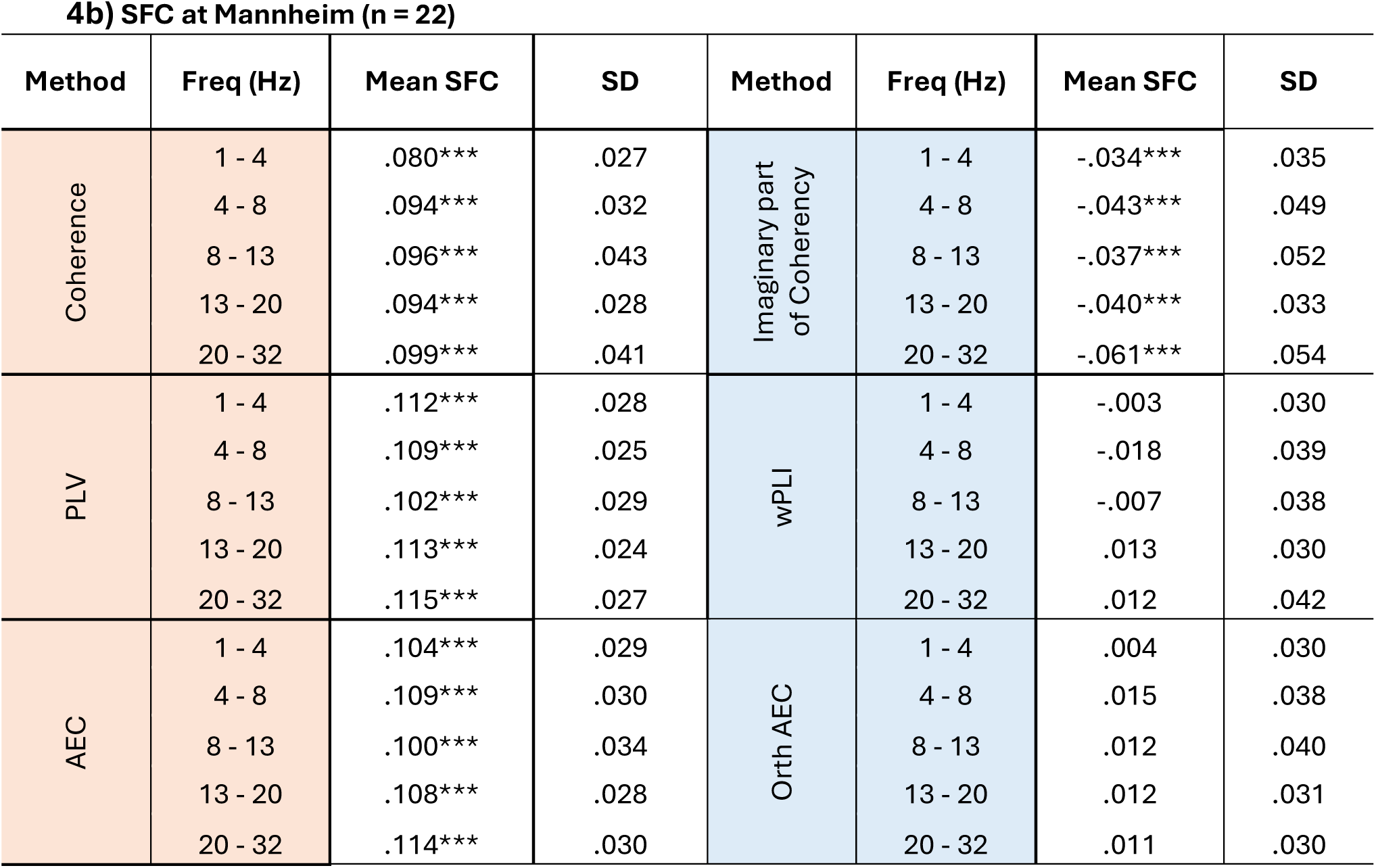
Structural-function concordance findings are replicated when analysing the **a)** KCL (n = 28) and **b)** Mannheim (n = 22) study sites separately. Across all frequency bands, concordance was significantly greater than 0 with zero-phase-inclusive methods (one-sided t-test or Wilcoxon sum-ranked test, *μ* = 0, *p* < .0001, Bonferroni corrected). Concordance was not significantly different from 0 for wPLI and orthogonalized AEC. Surprisingly, concordance was significantly less than zero for the imaginary part of coherency. Concordance for each zero-phase inclusive method was significantly higher than that for its methodologically corresponding zero-phase exclusive method (e.g. coherence versus imaginary part of coherency in 1-4 Hz), for each frequency band (paired samples t-test or Wilcoxon signed-rank test, *p* < .0001, Bonferroni corrected). *** concordance significantly different from zero, *p* < .0001, Bonferroni corrected. ^w^ = data not normally distributed, therefore one-sided Wilcoxon sign-rank test performed, otherwise one-sided t-tests performed. SFC = structure-function concordance; SD = standard deviation.

**Supplementary Table S5.**
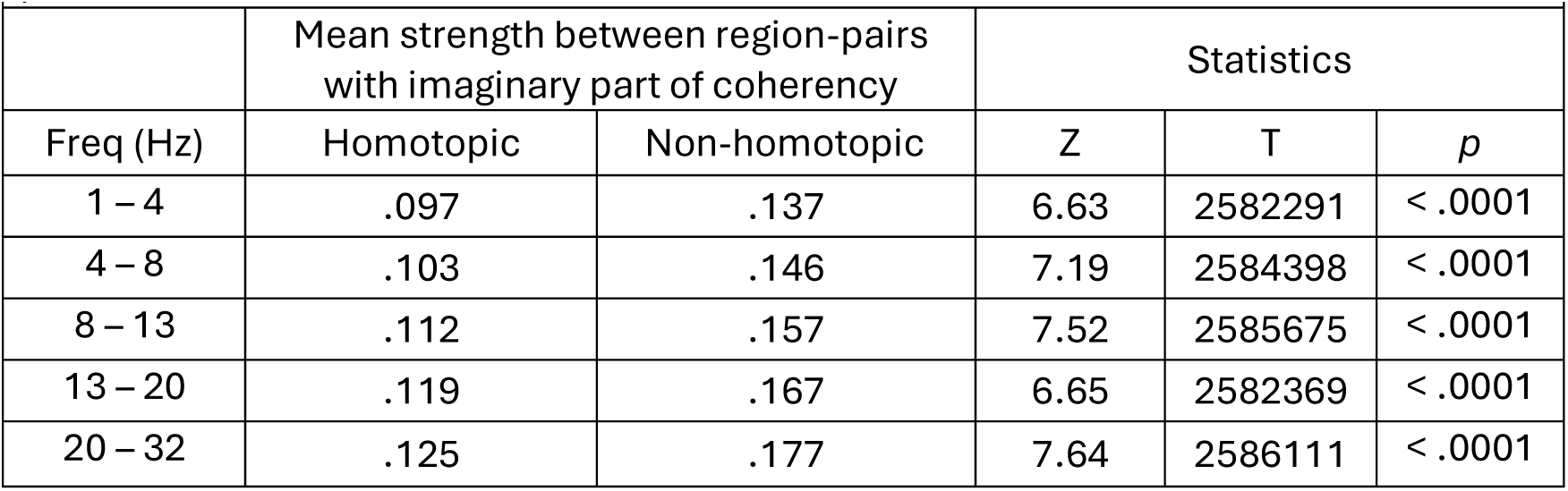
Comparison of mean functional connectivity strength between homotopic region-pairs versus all other region-pairs when using the imaginary part of coherency. Statistics calculated using the Wilcoxon rank sum test, *p* values Bonferroni corrected.

**Supplementary Table S6.**
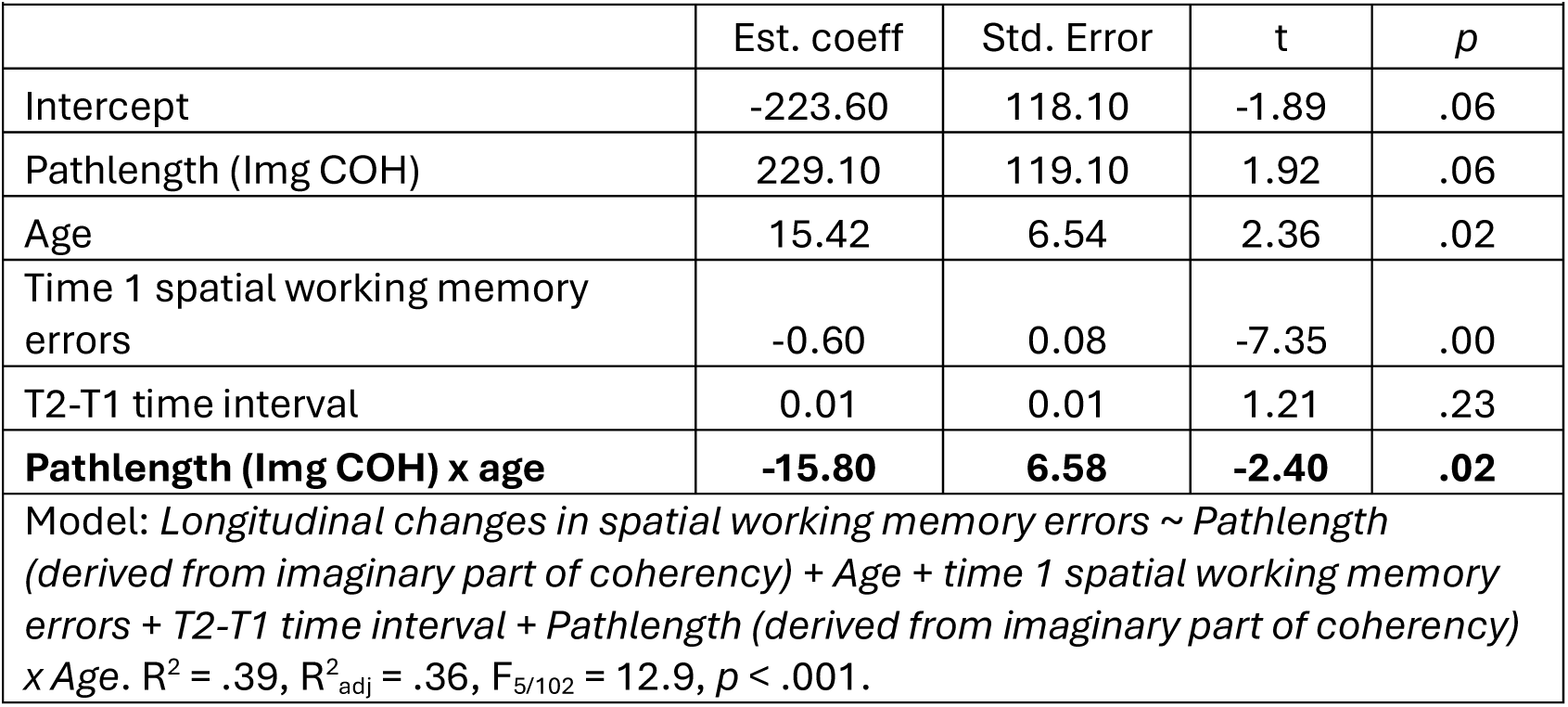
General linear model illustrating a significant age x pathlength interaction in predicting longitudinal changes in spatial working memory errors, when using the imaginary part of coherency. Participants aged 6-31-years.

**Supplementary Table S7.**
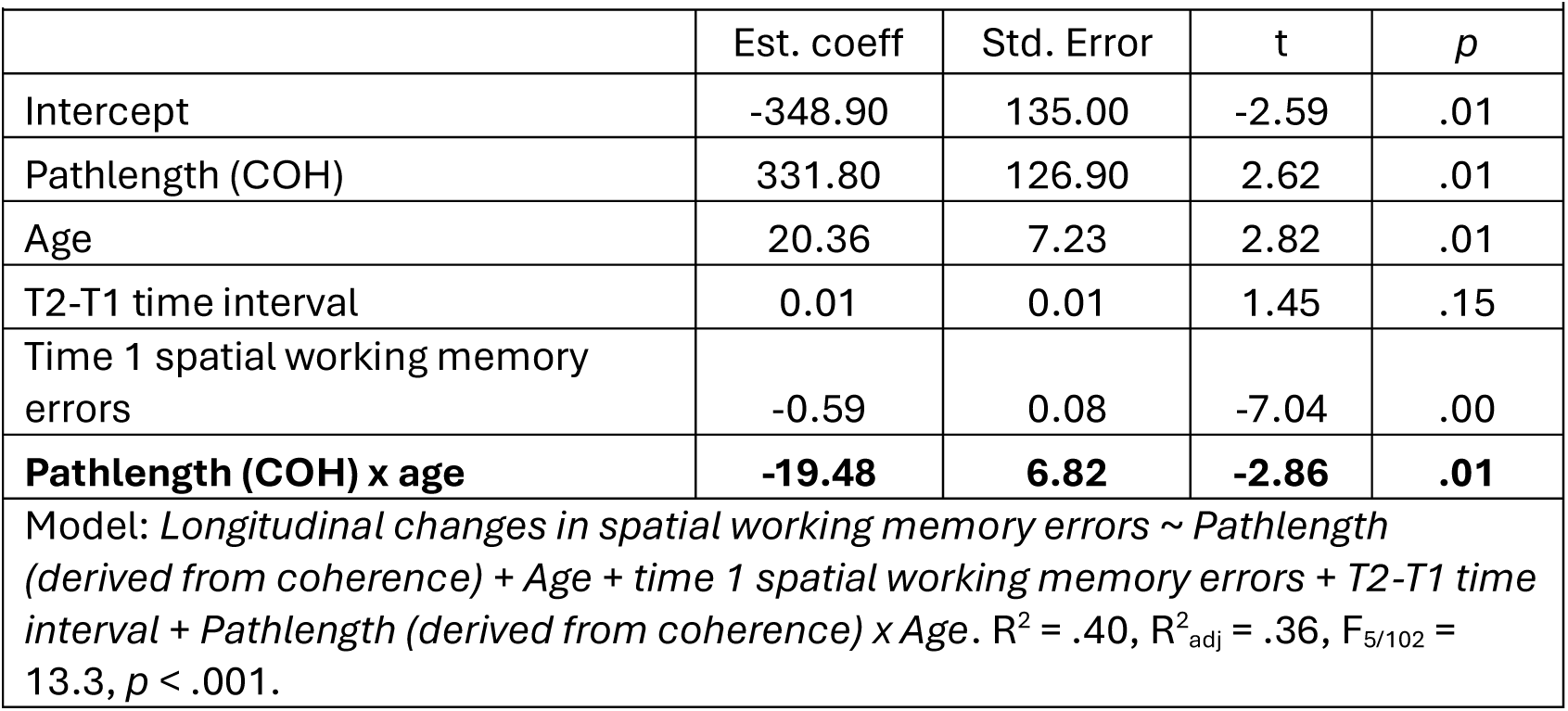
General linear model illustrating a significant age x pathlength interaction in predicting longitudinal changes in spatial working memory errors, when using coherence. Participants aged 6-31-years.

**Supplementary Table S8.**
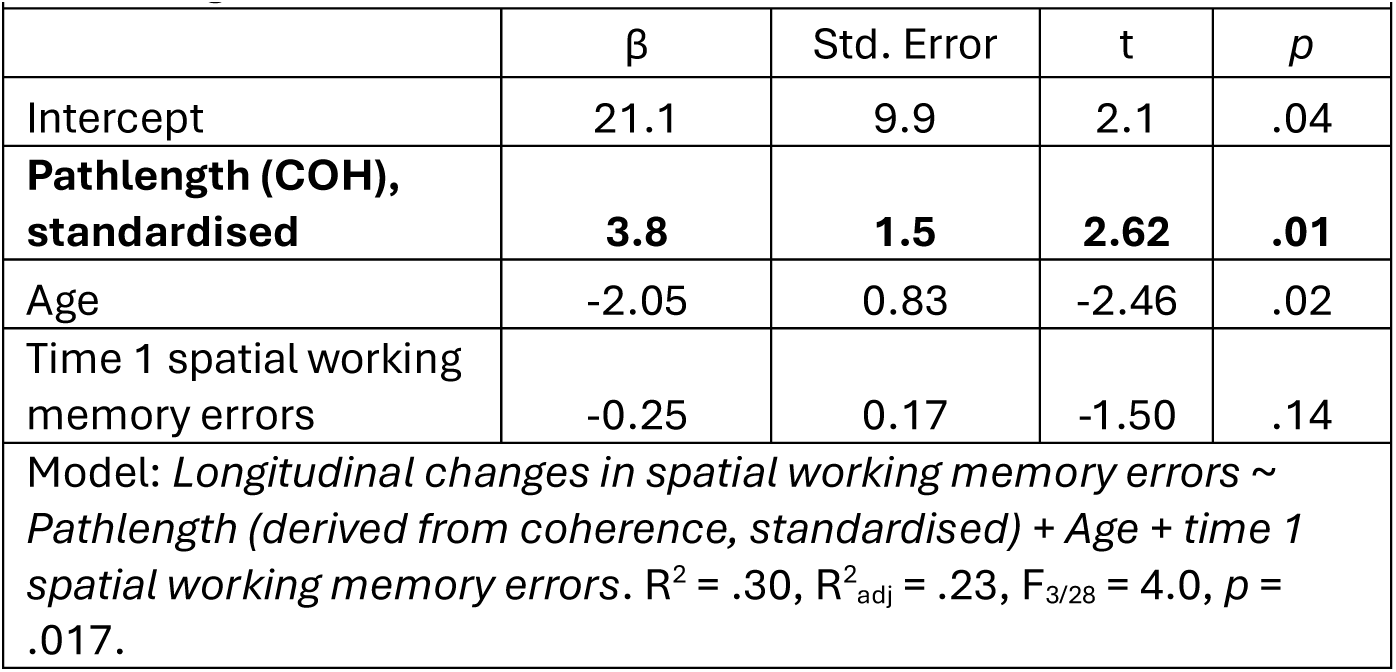
General linear model predicting longitudinal changes in spatial working memory ability in 6-12-year-olds using coherence.

**Supplementary Table S9.**
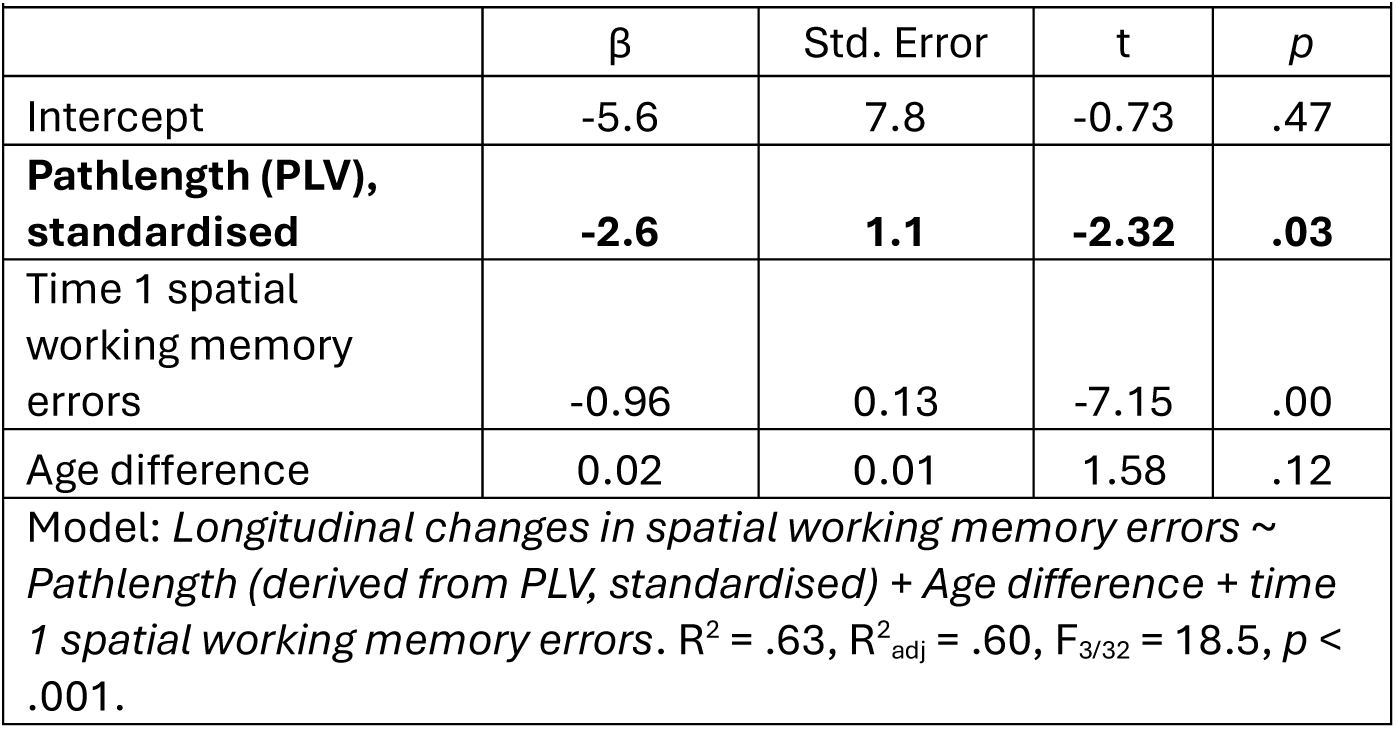
General linear model predicting longitudinal changes in spatial working memory ability in 18-31-year-olds using PLV.

