## Supplementary Figures and Tables for "Zero-phase-delay synchrony between interacting neural populations: implications for functional connectivity derived biomarkers"

### Supplementary Materials

**Supplementary Table S1 – Comparing the percentage of near-zero-phase-delay functional connectivity between homotopic- and heterotopic- interhemispheric region-pairs.** As there were 34 cortical regions per hemisphere, we randomly selected 34 heterotopic region-pairs, kept consistent across participants, to compare with the 34 homotopic region-pairs. DF = degrees of freedom.

| | Percentage of near-zero-phase-delay connectivity between interhemispheric regions | | Statistics | | |
|---|---|---|---|---|---|
| Freq (Hz) | Homotopic (%) | Heterotopic (%) | $X^2$ | DF | *p* |
| 1 – 4 | 84 | 55 | 6.05 | 1 | .014 |
| 4 – 8 | 86 | 58 | 5.44 | 1 | .020 |
| 8 – 13 | 83 | 50 | 8.19 | 1 | .004 |
| 13 – 20 | 87 | 61 | 4.57 | 1 | .033 |
| 20 – 32 | 89 | 67 | 3.10 | 1 | .078 |

**Supplementary Table S2 - Intrasession edgewise absolute agreement of EEG functional connectivity methods.** Median edgewise absolute agreement was calculated for each edge, using the interclass correlation coefficient (2,1; ICC). n = 99.

| Method | Freq (Hz) | Median ICC | IQR | Method | Freq (Hz) | Median ICC | IQR |
|---|---|---|---|---|---|---|---|
| Coherence | 1 - 4 | .15 | .26 | Imaginary part of Coherency | 1 - 4 | .02 | .16 |
| | 4 - 8 | .19 | .30 | | 4 - 8 | .03 | .15 |
| | 8 - 13 | .20 | .30 | | 8 - 13 | .07 | .16 |
| | 13 - 20 | .24 | .31 | | 13 - 20 | .03 | .16 |
| | 20 - 32 | .31 | .36 | | 20 - 32 | .07 | .18 |
| PLV | 1 - 4 | .69 | .29 | wPLI | 1 - 4 | -.01 | .13 |
| | 4 - 8 | .75 | .28 | | 4 - 8 | .02 | .14 |
| | 8 - 13 | .71 | .31 | | 8 - 13 | .17 | .18 |
| | 13 - 20 | .84 | .20 | | 13 - 20 | .04 | .15 |
| | 20 - 32 | .88 | .14 | | 20 - 32 | .09 | .17 |
| AEC | 1 - 4 | .37 | .43 | Orth AEC | 1 - 4 | .07 | .15 |
| | 4 - 8 | .47 | .45 | | 4 - 8 | .10 | .18 |
| | 8 - 13 | .48 | .42 | | 8 - 13 | .14 | .20 |
| | 13 - 20 | .57 | .42 | | 13 - 20 | .16 | .18 |
| | 20 - 32 | .71 | .36 | | 20 - 32 | .04 | .15 |

**Supplementary Table S3 – Intrasession consistency of the EEG functional connectivity adjacency matrix**. Zero-phase-inclusive methods had moderate to high consistency values, while zero-phase-exclusive methods had near 0 consistency values.

| Method | Freq (Hz) | Median Consistency | IQR | Method | Freq (Hz) | Median Consistency | IQR |
|---|---|---|---|---|---|---|---|
| Coherence | 1 - 4 | .34 | .16 | Imaginary part of Coherency | 1 - 4 | .07 | .09 |
| | 4 - 8 | .40 | .18 | | 4 - 8 | .10 | .11 |
| | 8 - 13 | .40 | .16 | | 8 - 13 | .12 | .11 |
| | 13 - 20 | .40 | .16 | | 13 - 20 | .11 | .12 |
| | 20 - 32 | .47 | .18 | | 20 - 32 | .16 | .16 |
| PLV | 1 - 4 | .80 | .10 | wPLI | 1 - 4 | -.01 | .07 |
| | 4 - 8 | .84 | .09 | | 4 - 8 | .01 | .11 |
| | 8 - 13 | .82 | .09 | | 8 - 13 | .12 | .16 |
| | 13 - 20 | .88 | .06 | | 13 - 20 | .02 | .08 |
| | 20 - 32 | .91 | .05 | | 20 - 32 | .02 | .08 |
| AEC | 1 - 4 | .55 | .15 | Orth AEC | 1 - 4 | .01 | .05 |
| | 4 - 8 | .64 | .18 | | 4 - 8 | .02 | .07 |
| | 8 - 13 | .60 | .15 | | 8 - 13 | .04 | .13 |
| | 13 - 20 | .66 | .14 | | 13 - 20 | .02 | .10 |
| | 20 - 32 | .71 | .13 | | 20 - 32 | .01 | .06 |

**Supplementary Table S4 - Structural-function concordance findings are replicated** when analysing the **a)** KCL (n = 28) and **b)** Mannheim (n = 22) study sites separately. Across all frequency bands, concordance was significantly greater than 0 with zero-phase-inclusive methods (one-sided t-test or Wilcoxon sum-ranked test, $\mu = 0$, $p < .0001$, Bonferroni corrected). Concordance was not significantly different from 0 for wPLI and orthogonalized AEC. Surprisingly, concordance was significantly less than zero for the imaginary part of coherency. Concordance for each zero-phase inclusive method was significantly higher than that for its methodologically corresponding zero-phase exclusive method (e.g. coherence versus imaginary part of coherency in 1-4 Hz), for each frequency band (paired samples t-test or Wilcoxon signed-rank test, $p < .0001$, Bonferroni corrected). *** concordance significantly different from zero, $p < .0001$, Bonferroni corrected. [w] = data not normally distributed, therefore one-sided Wilcoxon sign-rank test performed, otherwise one-sided t-tests performed. SFC = structure-function concordance; SD = standard deviation.

**a) SFC at KCL (n = 28)**

| Method | Freq (Hz) | Mean SFC | SD | Method | Freq (Hz) | Mean SFC | SD |
|---|---|---|---|---|---|---|---|
| Coherence | 1 - 4[w] | .112*** | .049 | Imaginary part of Coherency | 1 - 4[w] | -.041*** | .041 |
| | 4 - 8 | .119*** | .043 | | 4 - 8 | -.075*** | .042 |
| | 8 - 13 | .112*** | .035 | | 8 - 13 | -.057*** | .045 |
| | 13 - 20 | .116*** | .053 | | 13 - 20 | -.066*** | .060 |
| | 20 - 32 | .124*** | .050 | | 20 - 32 | -.081*** | .041 |
| PLV | 1 - 4 | .142*** | .040 | wPLI | 1 - 4 | .000 | .049 |
| | 4 - 8[w] | .142*** | .039 | | 4 - 8 | -.004 | .040 |
| | 8 - 13[w] | .132*** | .030 | | 8 - 13 | -.013 | .040 |
| | 13 - 20 | .144*** | .034 | | 13 - 20 | -.007 | .033 |
| | 20 - 32 | .146*** | .040 | | 20 - 32 | -.007 | .045 |
| AEC | 1 - 4[w] | .137*** | .046 | Orthogonalized AEC | 1 - 4 | .007 | .040 |
| | 4 - 8 | .133*** | .038 | | 4 - 8 | .006 | .035 |
| | 8 - 13 | .120*** | .047 | | 8 - 13 | .001 | .049 |
| | 13 - 20 | .140*** | .036 | | 13 - 20 | .006 | .039 |
| | 20 - 32 | .141*** | .039 | | 20 - 32 | .000 | .037 |

**b) SFC at Mannheim (n = 22)**

| Method | Freq (Hz) | Mean SFC | SD | Method | Freq (Hz) | Mean SFC | SD |
|---|---|---|---|---|---|---|---|
| Coherence | 1 - 4 | .080*** | .027 | Imaginary part of Coherency | 1 - 4 | -.034*** | .035 |
| | 4 - 8 | .094*** | .032 | | 4 - 8 | -.043*** | .049 |
| | 8 - 13 | .096*** | .043 | | 8 - 13 | -.037*** | .052 |
| | 13 - 20 | .094*** | .028 | | 13 - 20 | -.040*** | .033 |
| | 20 - 32 | .099*** | .041 | | 20 - 32 | -.061*** | .054 |
| PLV | 1 - 4 | .112*** | .028 | wPLI | 1 - 4 | -.003 | .030 |
| | 4 - 8 | .109*** | .025 | | 4 - 8 | -.018 | .039 |
| | 8 - 13 | .102*** | .029 | | 8 - 13 | -.007 | .038 |
| | 13 - 20 | .113*** | .024 | | 13 - 20 | .013 | .030 |
| | 20 - 32 | .115*** | .027 | | 20 - 32 | .012 | .042 |
| AEC | 1 - 4 | .104*** | .029 | Orth AEC | 1 - 4 | .004 | .030 |
| | 4 - 8 | .109*** | .030 | | 4 - 8 | .015 | .038 |
| | 8 - 13 | .100*** | .034 | | 8 - 13 | .012 | .040 |
| | 13 - 20 | .108*** | .028 | | 13 - 20 | .012 | .031 |
| | 20 - 32 | .114*** | .030 | | 20 - 32 | .011 | .030 |

**Supplementary Table S5 – Comparison of mean functional connectivity strength between homotopic region-pairs versus all other region-pairs when using the imaginary part of coherency.** Statistics calculated using the Wilcoxon rank sum test, *p* values Bonferroni corrected.

| | Mean strength between region-pairs with imaginary part of coherency | | Statistics | | |
|---|---|---|---|---|---|
| Freq (Hz) | Homotopic | Non-homotopic | Z | T | *p* |
| 1 – 4 | .097 | .137 | 6.63 | 2582291 | < .0001 |
| 4 – 8 | .103 | .146 | 7.19 | 2584398 | < .0001 |
| 8 – 13 | .112 | .157 | 7.52 | 2585675 | < .0001 |
| 13 – 20 | .119 | .167 | 6.65 | 2582369 | < .0001 |
| 20 – 32 | .125 | .177 | 7.64 | 2586111 | < .0001 |

| **Supplementary Table S6 –** General linear model illustrating a significant age x pathlength interaction in predicting longitudinal changes in spatial working memory errors, when using the imaginary part of coherency. Participants aged 6-31-years. | | | | |
|---|---|---|---|---|
| | Est. coeff | Std. Error | t | *p* |
| Intercept | -223.60 | 118.10 | -1.89 | .06 |
| Pathlength (Img COH) | 229.10 | 119.10 | 1.92 | .06 |
| Age | 15.42 | 6.54 | 2.36 | .02 |
| Time 1 spatial working memory errors | -0.60 | 0.08 | -7.35 | .00 |
| T2-T1 time interval | 0.01 | 0.01 | 1.21 | .23 |
| **Pathlength (Img COH) x age** | **-15.80** | **6.58** | **-2.40** | **.02** |
| Model: *Longitudinal changes in spatial working memory errors ~ Pathlength (derived from imaginary part of coherency) + Age + time 1 spatial working memory errors + T2-T1 time interval + Pathlength (derived from imaginary part of coherency) x Age*. $R^2 = .39$, $R^2_{adj} = .36$, $F_{5/102} = 12.9$, $p < .001$. | | | | |

| **Supplementary Table S7 –** General linear model illustrating a significant age x pathlength interaction in predicting longitudinal changes in spatial working memory errors, when using coherence. Participants aged 6-31-years | | | | |
|---|---|---|---|---|
| | Est. coeff | Std. Error | t | *p* |
| Intercept | -348.90 | 135.00 | -2.59 | .01 |
| Pathlength (COH) | 331.80 | 126.90 | 2.62 | .01 |
| Age | 20.36 | 7.23 | 2.82 | .01 |
| T2-T1 time interval | 0.01 | 0.01 | 1.45 | .15 |
| Time 1 spatial working memory errors | -0.59 | 0.08 | -7.04 | .00 |
| **Pathlength (COH) x age** | **-19.48** | **6.82** | **-2.86** | **.01** |
| Model: *Longitudinal changes in spatial working memory errors ~ Pathlength (derived from coherence) + Age + time 1 spatial working memory errors + T2-T1 time interval + Pathlength (derived from coherence) x Age*. $R^2 = .40$, $R^2_{adj} = .36$, $F_{5/102} = 13.3$, $p < .001$. | | | | |

| **Supplementary Table S8 –** General linear model predicting longitudinal changes in spatial working memory ability in 6-12-year-olds using coherence. | | | | |
|---|---|---|---|---|
| | β | Std. Error | t | *p* |
| Intercept | 21.1 | 9.9 | 2.1 | .04 |
| **Pathlength (COH), standardised** | **3.8** | **1.5** | **2.62** | **.01** |
| Age | -2.05 | 0.83 | -2.46 | .02 |
| Time 1 spatial working memory errors | -0.25 | 0.17 | -1.50 | .14 |
| Model: *Longitudinal changes in spatial working memory errors ~ Pathlength (derived from coherence, standardised) + Age + time 1 spatial working memory errors*. $R^2 = .30$, $R^2_{adj} = .23$, $F_{3/28} = 4.0$, $p = .017$. | | | | |

| **Supplementary Table S9 –** General linear model predicting longitudinal changes in spatial working memory ability in 18-31-year-olds using PLV. | | | | |
|---|---|---|---|---|
| | β | Std. Error | t | *p* |
| Intercept | -5.6 | 7.8 | -0.73 | .47 |
| **Pathlength (PLV), standardised** | **-2.6** | **1.1** | **-2.32** | **.03** |
| Time 1 spatial working memory errors | -0.96 | 0.13 | -7.15 | .00 |
| Age difference | 0.02 | 0.01 | 1.58 | .12 |
| Model: *Longitudinal changes in spatial working memory errors ~ Pathlength (derived from PLV, standardised) + Age difference + time 1 spatial working memory errors*. $R^2 = .63$, $R^2_{adj} = .60$, $F_{3/32} = 18.5$, $p < .001$. | | | | |
